# Modeling the mechanics of growing epithelia with a bilayer plate theory

**DOI:** 10.1101/2021.07.30.454446

**Authors:** Joseph Ackermann, Paul-Qiuyang Qu, Loïc LeGoff, Martine Ben Amar

## Abstract

Epithelia, which consists of cell sheets lying on a substrate, are prevalent structures of multi-cellular organisms. The physical basis of epithelial morphogenesis has been intensely investigated in recent years. However, as 2D mechanics focused most attention, we still lack a rigorous description of how the mechanical interactions between the cell layer and its substrate can lead to 3D distortions. This work provides a complete description of epithelial mechanics using the most straightforward model of an epithelium: a thin elastic bilayer. We first provide experimental evidence in *Drosophila* tissues that localized alterations of the cell-substrate (the extracellular matrix) can lead to profound 3D shape changes in epithelia. We then develop an analytical model modifying the Föppl-von Kármán equation with growth for bilayers. We provide a complete description of all contributions from biophysical characteristics of epithelia. We show how any localized inhomogeneity of stiffness or thickness drastically changes the bending process when the two layers grow differently. Comparison with finite-element simulations and experiments performed on *Drosophila* wing imaginal discs validate this approach for thin epithelia.

## 1 Introduction

Biological cell assemblies are often organized in laminar structures called epithelia. Epithelia cover most of our hollow organs, such as the esophagus [1], intestine [2,3] and stomach. The skin is also an epithelium constituted of several cell layers. The structure of epithelia is of one or several attached cell layers, which rest on a more disorganized polymeric substrate, called the extra-cellular matrix (ECM). Even the simplest epithelium, constituted of a single cell layer, must rest on an abutting ECM layer.

In the past decade, a lot of emphasis has been put on the role of the cytoskeletal cortex on the apical side of the cells (the side away from the ECM layer) in setting mechanical properties of epithelia [4]. However, recent investigations have established that the mechanical properties of the ECM may be as important in setting the shape of a tissue [5]. Both the cell layer and the ECM must be taken into account in the mechanical description of epithelia.

From the biomechanical viewpoint, an epithelium may be viewed as a bilayer made of two different soft materials with different stiffnesses and different ways to grow: the cell layer grows by mass increase and proliferation, while the ECM grows by addition of polymer-chains and swelling. A bilayer can also serve as a suitable approximation for more complex geometries, such as the multilayered skin, sub-divided into the dermis and the epidermis. In between layers of cells or in between cells and connective tissues, the ECM may develop into a stiff membrane enriched in collagen filaments called the basal membrane [6]. Epithelial cells may also have a reinforced cortex at their basal side, increasing the interfacial stiffness [7,8]. A more realistic mechanical view of epithelia would thus be that of a bilayer with the addition of stiff interfaces originating from the ECM or the cytoskeleton.

When one layer is a very soft substrate, several studies have attempted to deal with such bilayered systems [9–14] by treating the soft layer as an elastic foundation or as an ad-hoc resistance potential [10,11]. These models become essentially single-layered and cannot account for the diversity of behavior of two-layered systems where the two layers play similar roles and have similar properties. We aim here to present a simple mechanical model of thin bilayers using the Föppl-von Kármán equations (FvK) with growth. In the context of a thin plate hypothesis, the FvK equations allow a 3D to 2D dimensional reduction which considerably simplifies the analytical treatment of plate mechanics. The FvK equations are thus well adapted to incorporate many different biological features of layered tissues which are difficult to handle analytically in a fully three-dimensional formalism. It has recently been shown that FvK equations with growth allow to explain the buckling of thin objects such as flowers and algae in a rather simple way [15,16], when compared to the full treatment of finite elasticity with growth [17]. In addition, a slight modification of these equations allows to treat initially weakly curved membranes. This prompted us to use the same approach for a bilayer with growth. When modeling a single-layered epithelium, the two layers of the model are the ECM and the cells, while the interfacial stiffness represents the basal cytoskeletal cortex of cells (left inset in fig. (1)). When modeling a more complex system, such as the skin, one layer of the model represents the cell layers of the epidermis, the second layer is the dermis (a connective tissue) and the interfacial stiffness represents the extra-cellular basal lamina (right inset in fig. (1)), see reference [6]. Our model can account for local variations in the stiffness or in the thickness of the layers as well as interfacial stiffnesses. Such variations are often present in biological tissues, and are thought to be shape generators in morphogenetic processes. These local variations in mechanical properties can also be induced in the context of perturbative experiments.

**Fig. 1.**
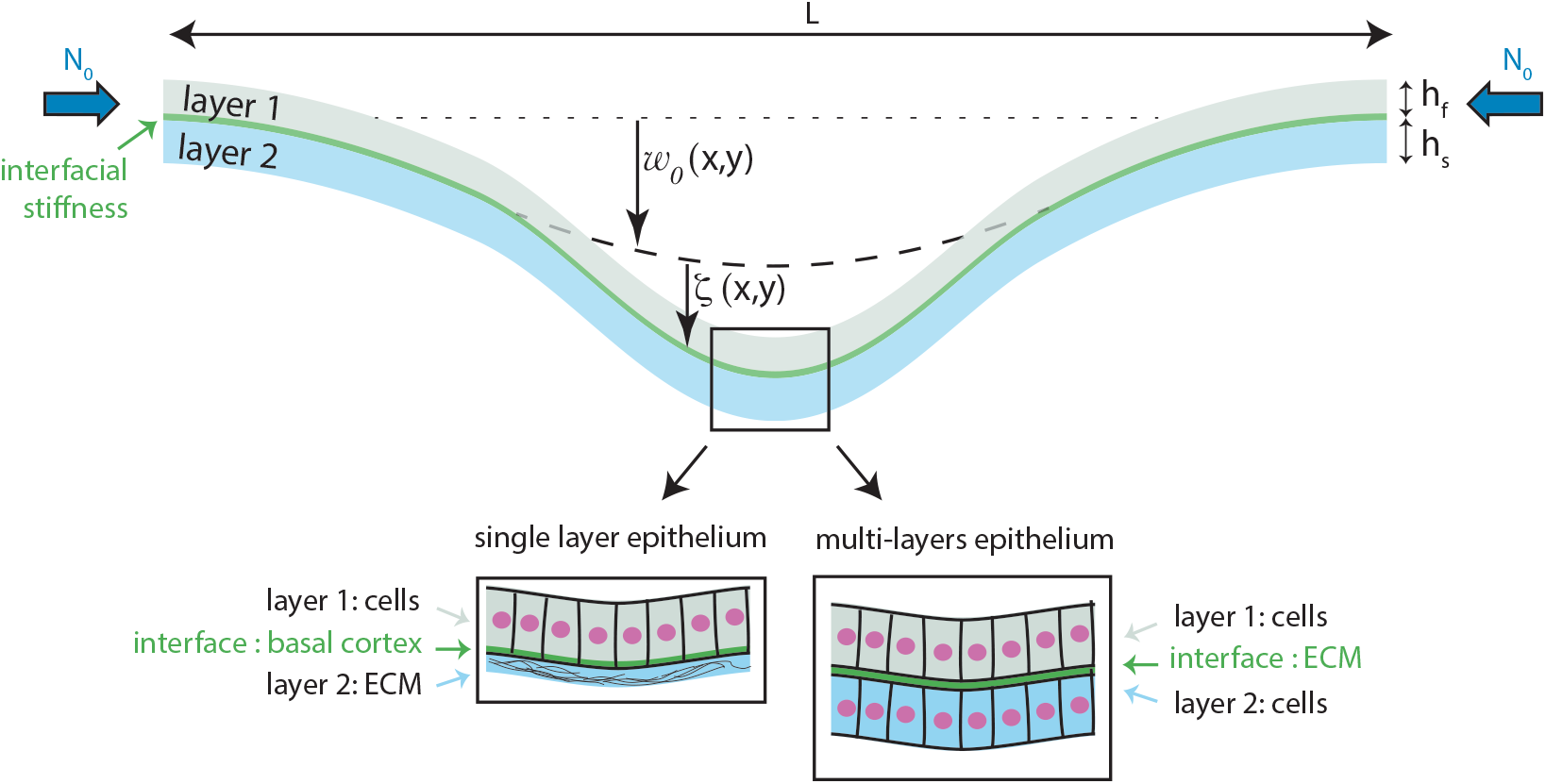
Geometry of the bilayer model. *w*0(*x, y*) is the deviation of the initial configuration from a flat surface; *ζ*(*x, y*) is the displacement from this initial configuration as a result of elastic energy minimization under the action of growth and pre-stress (*N*0). The schema at bottom represents biological context that can be modeled by the theory: the two layers can either be cell or ECM layers of different height, Young modulus and growth rate; the stiff interface can be a cytoskeletal cortex or a thin ECM.

As much as elasto-mechanical processes may drive the shape of epithelia, this shape builds upon a long and intricate history of growth, stress distribution, and changes in mechanical properties. On the other hand, our formalism, which uses linear elasticity with moderate non-linear elastic strains, can only account for slight shape variations, spanning time intervals typically smaller than the doubling time of cells within the tissue. To mitigate this problem, we encapsulate all the previous history of development and morphogenesis in an initial shape and a pre-stress. This allows confrontation with experiments where the tissue, at the onset of observation, is rarely in a flat and stressless configuration and where the full history of development is also inaccessible. We show that the model can treat many aspects of biological tissues, which are not so common in material sciences. Although the formalism was initially developed to treat biological growth, it could also serve for the opposite case of resorption, which could be of interest both in a biological and material science context.

In this paper, we first demonstrate experimentally how alteration of the ECM can substantially impact the shape of an epithelium. For this, we genetically degrade the ECM in a band of cells in the Drosophila wing imaginal disc -the precursor of the adult wing. Subsequent sections are devoted to developing an analytical and numerical treatment of a bilayer model. We aim to account for the experimentally observed distortion of the tissue and provide a general framework to address the mechanics of growing bilayered tissues at large. Section (3), is devoted to the geometry of the sample under study and a reminder of the necessary approximations to incorporate the formalism of finite elasticity with growth into the FvK approach. In sect. (4), we give the main equations for a bilayer and an approach of the treatment of the interface. In sect. (5), we derive the Euler-Lagrange equations for the elastic bilayer equivalent to the FvK equations and we demonstrate the peculiar role of the neutral surface for the bending of the plate. Its position is derived for arbitrary stiffness and thickness, not necessarily constant, of the two layers. Section (6) focuses on uniaxial deformations, which simplify the FvK modified equations for a bilayer with additive terms at the origin of buckling. In this section, numerical results illustrate the various cases for pre-stressed plates but also for slightly curved membranes, and the theory is confronted with experimental data. In sect. (7), results obtained with finite element simulations achieved in the same context of growth, thickness, stiffness and defects are presented. Finally, we conclude in sect. (8) by giving some perspectives.

The model can be adapted to many other biological systems, as epithelia are ubiquitous in most living species. It could also be used for other thin objects such as leaves or algae.

## 2 Experimental motivation: biomechanics of *Drosophila* wing imaginal discs

The Drosophila wing imaginal discs (the precursors of the adult wing) are epithelial tissues that became, over the years, one of the most studied and best characterized systems to study growth [18]. The growing wing imaginal disc displays a highly patterned field of mechanical stresses [19–21]. Cells at the periphery of the epithelium sustain a strong mechanical stretch. This pre-stress builds up as the tissue grows. In addition, the tissue becomes curved as development proceeds, gradually changing from a simple flat surface to a more complex curved surface.

Until recently, most attention was brought upon the role of the apical surface of the epithelium in setting its mechanical properties. It was however recently demonstrated that the extracellular matrix upon which the epithelium sits plays a major role in shaping the tissue. Alterations in the basement membrane composition, which includes collagen IV, laminin, nidogen, and heparan-sulfate proteoglycans, have profound effects on the shape of the wing imaginal disc [22]. Alterations of the basement membrane have also been shown to play an active part in setting the 3D shape of the wing disc by promoting the formation of the folds that gradually arise in the tissue [23,24]. From these experiments, it now stands that wing imaginal discs, and most epithelia in general, are two layered composite structures. Both the cell layer and the ECM layer take an active part in shaping the tissue [25].

To assess the role of basement membranes in setting mechanical properties of the imaginal discs, we used *Drosophila* genetics to express the matrix-cleaving metallo-protease Mmp2 in a band of cells, located at the center of the wing imaginal disc along the Dpp-Gal4 genetic pattern (fig. (2a)). The genotype of the observed tissues is ubi-cad: GFP UAS-GFP tub-Gal80_ts_/+; Dpp-Gal4/UAS-Mmp2. It combines the necessary transgenes to express the metallo-protease locally and to visualize cells with a cadherin: GFP fusion. The biochemical action of Mmp2 is to degrade structural components of the ECM [26], which in mechanical terms implies a possible change of ECM stiffness as well as the thickness. We cannot rule out also an indirect effect on the basal cytoskeletal cortex of cells, as an altered ECM in the vicinity of the cell layer implies a reduced activation of integrin receptors. Our elastic model will need to take into account these different contributions. The metallo-protease expression is controlled in time via the thermo-sensitive mutant of Gal80, by switching *Drosophila* from 18°C to 29°C 24h before observations (fig. (2b)). Such a timing corresponds to approximately 18 hours of Mmp2 expression since 6 hours are required to reach full Gal4 expression after the temperature switch [27]. To image the imaginal discs, we performed *ex-vivo* cultures of wing imaginal discs as in [19]. The living tissues were then imaged with a spinning disc confocal microscope.

**Fig. 2.**
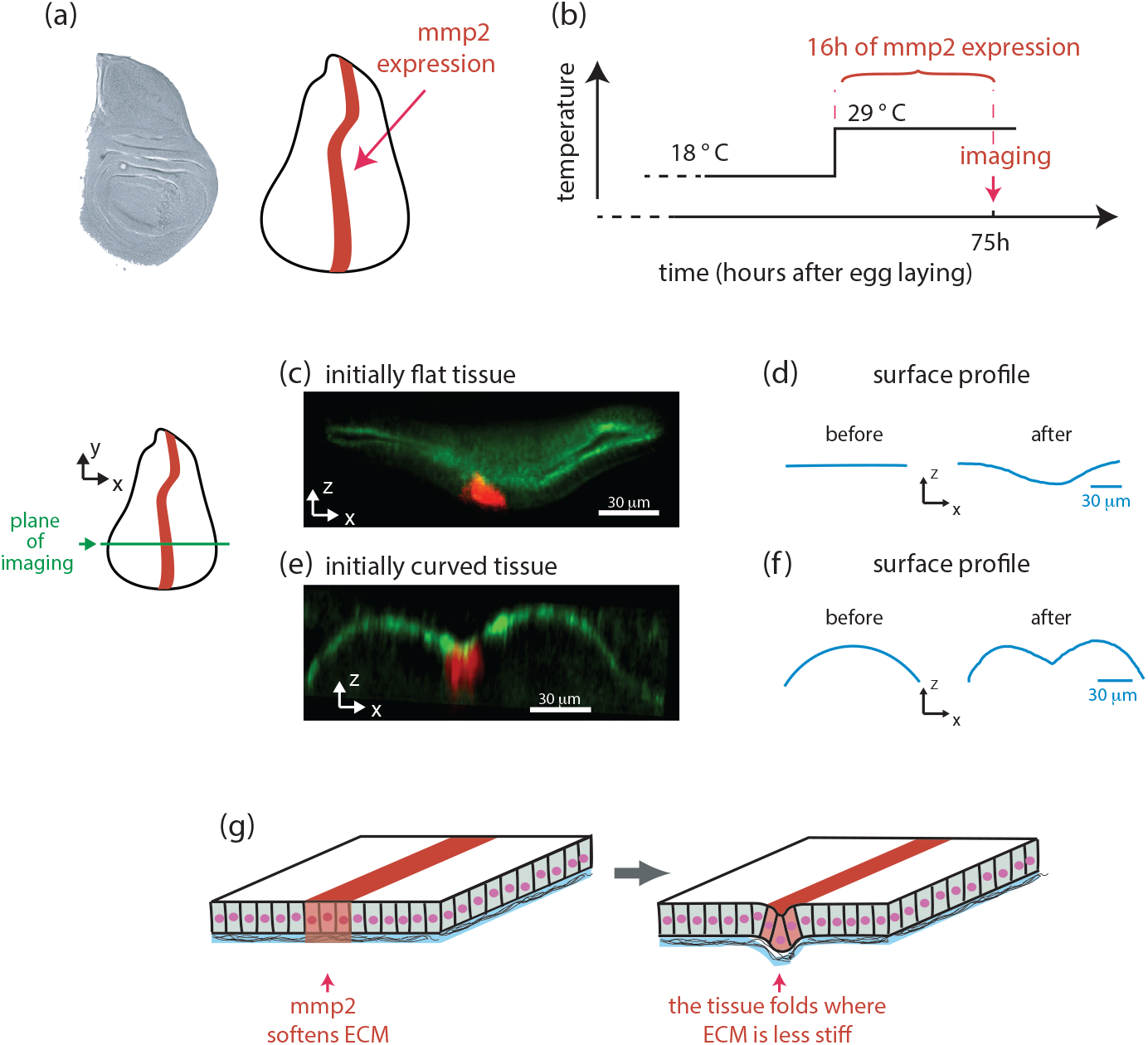
Folding an epithelium by local ECM degradation. (A) Transmission light imaging of a *Drosophila* wing imaginal disc (left) and corresponding sketch with the region of localised expression of the protease Mmp2 shaded in red (right). (B) To control expression of the metalloprotease, the sample is switched from 18°C to 29° C several hours before observation. (C-F) Experimental demonstration. (C) An xz cross section demonstrates the shape of an initially flat wing imaginal disc under the effect of a localized metalloprotease expression (in the red colored region). The schema in (D) demonstrates the change in shape of the tissue. (E,F) Same as (C,D) but with an initially curved wing imaginal disc. (G) A schematic representation of the shape changes observed in the wing imaginal disc.

Figure (2c) shows the cross-section of an imaginal disc 18 hours after metalloprotease expression. The region of perturbation, identified by RFP expression, is shaded in red on the figure. Figure (2d) shows the profile of the apical surface of the epithelium, which has a region of inflection near the protease expression. This profile (fig. (2c,d)) corresponds to a wing imaginal disc that was initially flat. Alternatively, fig. (2d,e). shows the same perturbation outcome on a wing imaginal disc that was initially curved - curvature normally builds up at late stages of wing disc development (around 80 hours after egg laying). The resulting tissue is very different from the previous case. This time, the profile shows a more complex configuration: the naturally occurring curvature is prevalent on the borders of the wing disc; at the same time, the metalloprotease action in the region of perturbation induces an inversion of the curvature there.

To conclude, we observed that upon ECM degradation (which may impact the stiffness, the thickness, and the interfacial mechanics), the epithelium curves in the perturbation region (fig.(2h)). We observed two archetypal outcomes depending on whether the tissue was initially flat or curved, which can lead to the inversion of the naturally occurring curvature in the perturbation region. These experimental observations will be subsequently confronted to our FvK bilayer model and our finite-element simulations (see fig. (8) and fig. (10)).

## 3 Modeling

The geometry of the model is represented in fig. (1): two layers of different thickness (*h*_*f*_ and *h*_*s*_), stiffness and growth rate are organized in a bilayer, with an interface which may also bear a stiffness. The shape of the initial configuration (see explanations in fig.(1) is given by w_0_(*x, y*), the deviation from the horizontal line; the vertical deflection arising from the elastic minimization is denoted *ζ*(*x, y*). There are no explicit rules about the difference in stiffness or thickness of the two layers, but they tend to be of the same order of magnitude for biological systems. While there are also no rules regarding the thin interface layer, it will only play a role if it is much stiffer. Contrary to inert materials, changes in the shape of the structure arise from inner processes such as volumetric growth rather than through external loading, although this latter is not eliminated *a priori* from the model. The bilayer can model, with different degrees of accuracy, a single or a multi-layered epithelium.

A decade ago, Dervaux *et al*. established the formalism of a growing plate using the theory of finite elasticity with growth [15,16]. Here, we extend their analysis of a single layer to a thin bilayer, both layers having different growth rates and different elastic properties but remaining thin. Indeed, the FvK equations rest on some limitations: the order of magnitude of the vertical deflection *ζ* due to the buckling and the initial deviation from the horizontal line w_0_ must remain small compared to the horizontal length L (they may be larger than the thickness h of the plate).

In the following, we first present the model for the general 2D case with arbitrary buckling deformations, and subsequently treat the simpler uni-axial folding.

### 3.1 The geometric and elastic strain

We consider two different layers with thickness h_f_ (top) and h_s_ (bottom). When volumetric growth or resorption occurs in an elastic sample, each point is displaced and the gradient of this displacement is a geometric tensor called the deformation tensor: ***F***. According to the famous *Kroner-Lee* decomposition [28], ***F*** results from both the elastic tensor ***A*** and the growth tensor **G** in a simple way :

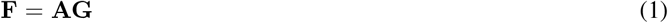

Here, plasticity is neglected. The amount of growth per unit volume is given by Det ***G*** − 1, which is negative for mass resorption and positive for growth. When it is small, the growth tensor **G** can be decomposed into : 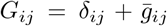 with 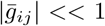 In addition, in the case where the external load maintains a low level of strain, the geometric deformation tensor is then also small leading to the following displacement of the points originally located at ***r*** = (*x, y, Z*) :

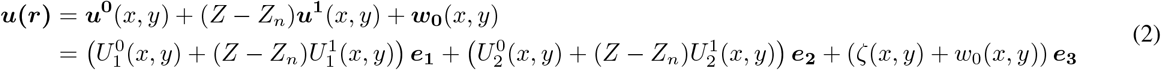

where *w*_0_(*x, y*) represents the initial position of the interface bilayer, and *ζ*(*x, y*) the deflection due to the buckling event. Each **e**_**i**_ represents a cartesian unit vector and the superscripts “0” or “1” stand for the order of perturbation. Indeed, the ratio between the thickness of the bilayer and the horizontal size *L* is one small parameter ϵ_1_ = h/L but the magnitude of the displacement in the *Z* direction compared to *L* is a second independent small parameter ϵ_2_ = *ζ*/L. We assume that both layers have comparable thickness and h means either *h*_*f*_ or *h*_*s*_. In the classical FvK approach, 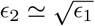 a scaling which is justified below and which will constrain 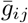 and w_0_. Since we examine a composite material here, the choice of the origin of the *Z* axis is free at this stage. We define this origin at the physical interface between both layers. We also introduce *Z*_*n*_, a possible shift in the expansion of order *h*. This surface defines the surface of separation between parallel surfaces in extension compared to the ones in compression due only to the buckling process. In the single plate geometry, *Z*_*n*_ corresponds to the position of the neutral surface and is located at *h*/2. In the following, an analytical expression for *Z*_*n*_ will be derived. We can then define the geometric deformation gradient :

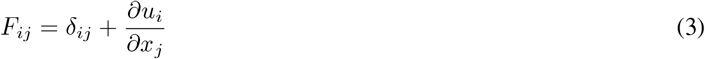

where, as in Landau and Lifchitz’s book of elasticity [29], the index *i* indicates an index varying between 1 and 3 while greek letters restrict to 1 or 2. The elastic strain that we deduce from eq.((1)) when A and G are close to unity reads :

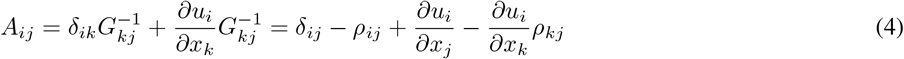

where we have defined the tensor ***ρ*** such that ***G***^***−1***^ = ***I − ρ***, each component of *ρ* remaining a small quantity to validate the FvK approach. *ρ* is not rigorously equal to (***G* − *I***). They are equivalent only at linear order. In addition, we assume no change of the thickness of the sample which means *ρ*_33_ = 0, this hypothesis simplifies the equations and can be easily revisited. A spatially constant value of *ρ*_33_ over the sample simply enters in the definition of *h*. We also assume that *ρ*_*α,β*_ scales as 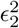 while *ρ*_*α,β*_ and *ρ*_3α_ scale as ϵ_2_. We also introduce a new tensor **g** whose components are:

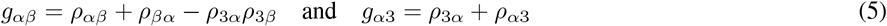

This tensor is symmetric and has the same scaling behavior as *ρ*. It will simplify equations later on. In the general case, the growth tensor **g** is a function of the coordinates, including *Z*. This is of interest in the context of epithelia, as the cells may not grow in the same way as the ECM substrate. A discontinuity may therefore develop at the interface. Similarly, the structural material coefficients such as the Young modulus will be different in both layers. For simplicity, we do not expand these coefficients and we keep their full variations. We expand only the strain and stress tensors in power of *Z* -*Z*_*n*_. This assumption is indeed perfectly exact if the bilayer is made of two perfect homogeneous layers. To order 2 in ϵ_2_, the elastic strain ϵ reads:

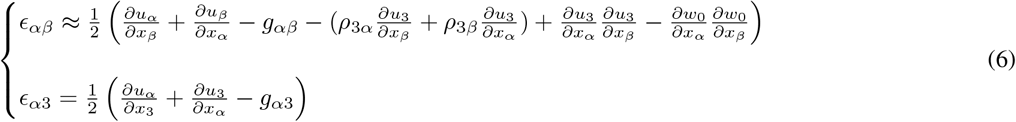

where we only keep the quadratic term in *u*_3_ = *ζ* + *w*_0_(*x, y*) for the deformation tensor, all the other contributions being neglected. Such simplification is justified by the choice of the traditional scalings of the FvK equations which favours the bending contribution, of order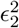, compared to the in-plane terms of order ϵ^2^. It is clear that any change in the scaling approximation for the strains as done for the enhanced FvK or Koiter-Sanders model [30] will complicate a lot the equations in the presence of growth. We will not treat this in the present work. For a full analysis of the scalings in initially curved plates, see Chap. 6 of [31]). Our formulation is slightly different from the FvK-Donnell formulation [10,32]. but gives similar results after the variational process of the elastic energy so after integration by part. It recovers perfectly the results of Ciarlet and Miara [33]. The last terms in eq.(6) represent the geometric stretch of the deformation in the *x*_3_ direction.

For a consistent expansion, all terms in this relation must have the same order of magnitude. This implies several scaling relations. We get for the horizontal deformation: *u* ≃ *ζ*^2^/*L*, for the growth element *ρ*_*αβ*_ ≃ *ζ*^2^/*L*^2^ and for the representation of the initial stress-free configuration, *w*_0_ ∼ *ζ*.

### 3.2 The monolayer case

We aim here to establish the equilibrium equations of a growing bilayer under initial lateral loading represented by *N*_0_ in fig. (1), when it is initially planar or weakly curved. To this end, as in Landau and Lifshitz book [29] and in Refs [11,15,16], our strategy consists in calculating the elastic energy and deriving the Euler-Lagrange equations through variations. Most importantly, we write the stress and strain tensors as a function of *x* and *y* only. First, we consider each layer independently since each of them has its own elastic coefficients and its own growth characteristics. Due to the weakness of the deformation for slender objects, we can apply the Hooke’s Law which is the constitutive equation of linear elasticity, but we maintain the non-linearities of the strains according to the strategy of the FvK equations. We also consider the usual membrane hypothesis: *σ*_*i*3_ =0 which is equivalent to the plane stress approach. Then, it reads for the Cauchy stress components *σ*_*αβ*_ where *α* and *β* are restricted to *x, y*:

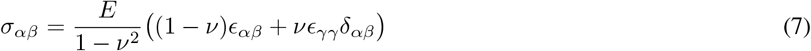

with *ν* ≃ 1/2 for the incompressible case, an assumption commonly used for living tissues. In order to determine the elastic energy *ϒ*, we then write the elastic strain tensor (also a function of *x* and *y*), which we decompose in orders of (*Z* − *Z*_*n*_) with the help of the expansion 2), so that

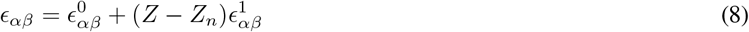

The zeroth order reads:

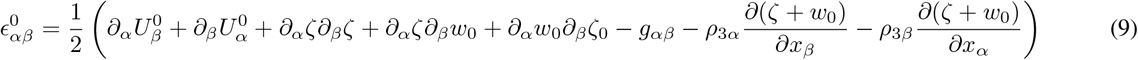

The first order results from the cancellation of the stresses of the third dimension : σ_33_ = σ_13_ = σ_23_ = 0, so ϵ_13_ = ϵ_23_ =0 (see [29]) and using the definition eq.(6), we derive:

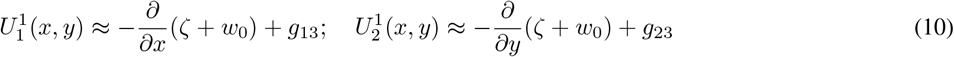

Thus the strains at first order become :

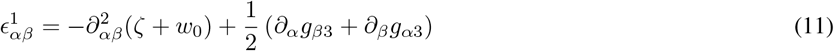

Following the equilibrium equation eq.(7) and the decomposition of the elastic strain (eq. (9) and eq. (11)), we can also decompose σ_*αβ*_ as 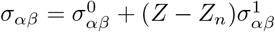.

## 4 FvK equations for the bilayer case

As in the presentation achieved in Landau and Lifchitz’s book of linear elasticity [29], we average the elastic energy in the thickness of the bilayer.

### 4.1 Averaging the FvK equation

An important problem in the construction of the model is to establish the position of the neutral surface when the layer is not perfectly homogeneous or for multiple layers. According to Ref [29], the neutral surface defines the separation between layers in compression from layers in tension when the structure is weakly bent. For an homogeneous sample, the neutral surface is naturally the middle surface of the sample. But for a bilayer, the position of the neutral surface is unknown. The interface between both layers seems to be the natural choice for the definition of the origin of the *Z* coordinate but it is not the neutral surface. So we focus first on its position, then we proceed to the process of variation of elastic energy.

#### 4.1.1 Position of the neutral surface

Despite the fact that attention was given to this question for functionally graded beams [34,35,30,36], the determination of *Z*_*n*_ has not been found in the literature. Here, we propose a simple argument and we suggest decomposing the full energy into:

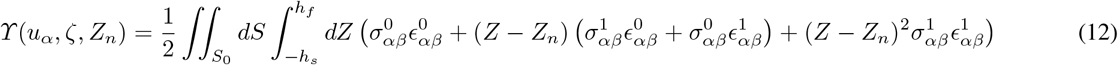

Since *ϒ*(*u*_*α*_, *ζ, Z*_*n*_) must remain positive independently of the growth coefficients and the strains, this happens automatically if the intermediate term cancels then imposing:

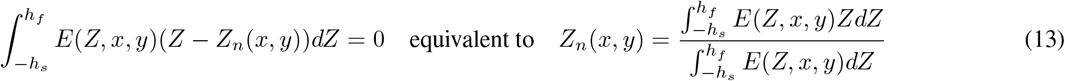

average position of the bilayer weighted by the Young modulus. In the general case, there is no guarantee that the neutral surface *Z*_*n*_ is positioned inside the bilayer. However if in each layer the Young modulus is a constant (but different from top to bottom), the neutral surface is located inside the bilayer.

Finally, we derive:

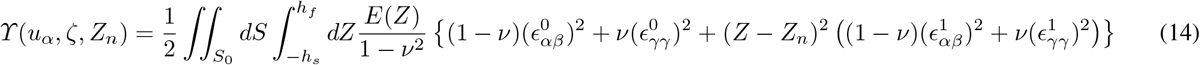

This method has been employed by Kozlov and Winterhalter [37] for the determination of the neutral surface of strongly curved lipidic membranes.

#### 4.1.2 Variational determination of the Euler-Lagrange equation

The Euler-Lagrange equations result from variations of ϒ with respect to *ζ* and *u*_*α*_, the linearity between **σ** and **ϵ** leading to *δ⥾* = ∫ *d**r**σ*_*αβ*_*δϵ*_*αβ*_ These variations, taken one after the other, must vanish at linear order. Special attention must be given if the growth tensor or the Young modulus *E* are dependent on *Z* as it is obviously the case for the bilayer, and sometimes of the other coordinates *x* and *y*. Coming back to the definition of *ϒ* as:

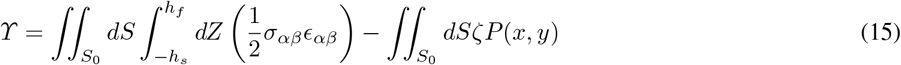

where *P* is a vertical external loading. The variation of ⥾ with respect to the horizontal deformation *uα* is easily derived and yields:

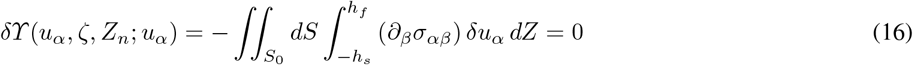

which gives that *∂*_*β*_*σ*_*αβ*_ = 0 everywhere in the sample. This corresponds to the second equation in the FvK formalism [29], sometimes re-written with the Airy potential, which is very useful for analytical or numerical solutions of a 2D problem. Variation of *ϒ* with respect to *ζ* gives:

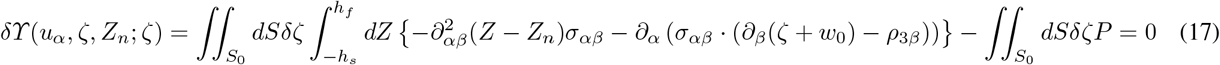

In eq.(16,17), the two bracketed terms must vanish. We do not mention explicitly the contributions coming from integration by part which fixes the boundary conditions for free boundaries. In the following, we consider only clamped boundary conditions.

#### 4.1.3 The averaged Euler-Lagrange equation for the growing bilayer

It is now possible to transform eq.(17) once we define the new bending coefficient and the new spontaneous curvature due to growth:

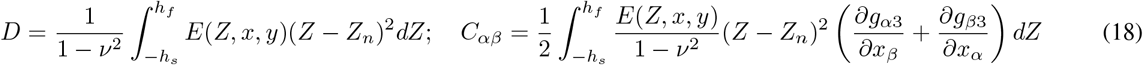

Doing again the separation of σ into σ^0^ and σ^1^, an intermediate step consists in integrating σ_*αβ*_ over the *Z* variable, which gives first:

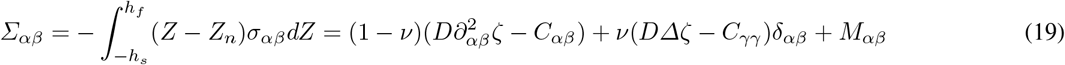

where we have dropped the dependence of *D* and C_*αβ*_ in *x, y* to simplify notations, introduced the 2D Laplacian Δ = *∂*_*αα*_, and a new tensor *M*_*α,β*_:

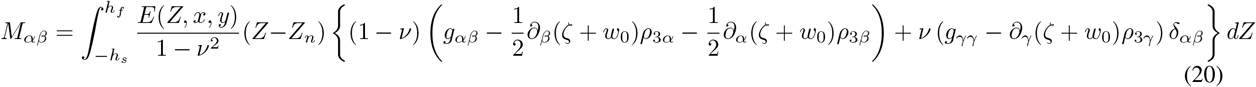

The simplification of *M* is due to the definition of the neutral surface and the fact that only the growth component *g*_αβ_ may depend on *Z*. If it is not the case, *M*_*α β*_ vanishes. At last, eq.(17) gives the equivalent of the first FvK equation:

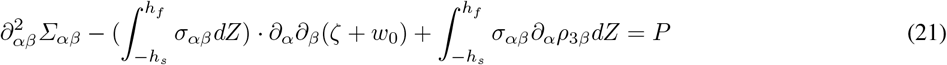

This last equation, eq.(21), governs mostly the bending deformation. In the nonlinear regime of the FvK formalism, it is coupled to the horizontal stresses σ_*α,β*_ whose equilibrium is given by eq.(16). When not stated otherwise, σ_*αβ*_ is the total stress. *D* appears to be an effective bending coefficient of the composite structure and *C*_*αα*_ as a spontaneous curvature associated with growth. For a homogeneous layer with no growth *Z*_*n*_ = *h*/2, *M*_*α,β*_ vanishes and one recovers the traditional FvK equations without growth, see [29]. Notice that in eq.(21), we also recover the spontaneous curvature κ = *∂*_*α*_*g*_*α*3_ found in [16] and the curvatures, inverse of the radii of curvature, of the initial shape given by the tensor *∂*_*α,β*_*w*_0_. Let us focus now on the uniaxial folding where all the equations simplify.

### 4.2 FvK equations governing uniaxial folding

For uniaxial folding, the only dependence is along the x axis and all the tensors are reduced to one element, which simplifies a lot the analysis but reduces the diversity of observed buckling patterns, as explored in previous works [12,14]. The elastic plane stress equilibrium equation, eq.(16) gives ∂_*x*_σ _11_ =0 so it is only a function of *Z* and *N* defined by:

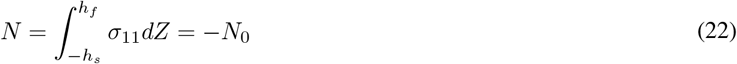

*N*_0_ indicates a lateral integrated compression, a situation easily realized in material sciences but in embryos it results from accumulated pre-stress originated from the imaginal disc and the connected tissues at the boundaries [38,20] (see fig.(1)). The bending equilibrium equation now reads:

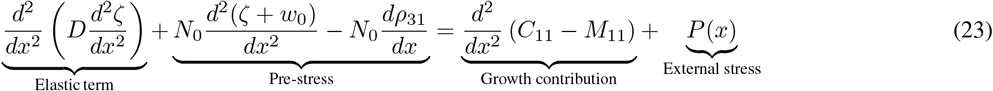

where we have neglected the *Z* dependence of *ρ*_31_ involved in eq.(21). We detail *M*_11_ in the simplified case of growth occurring only in a single layer.

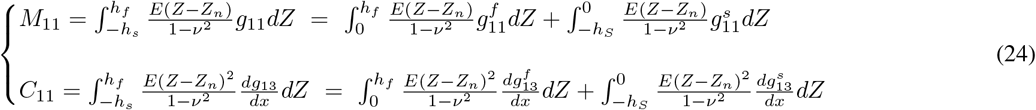

We have derived an effective equation for the bilayer, which takes into account that the growth may differ from the top to the bottom layer via *C*_11_ and *M*_11_. Contrary to *C*_11_, *M*_11_ appears only if the growth tensor component *g*_11_ is different in both layers. If in addition, the second derivative of these two quantities does not vanish, they appear equivalent to a vertical loading acting on the bilayer giving it a curvature in absence of an initial one (*w*_0_ = 0). This will destroy the perfect symmetry up and down, once averaged over the thickness of the bilayer and explain the bending in the case of growth. We now examine in more detail how the structure of the bilayer and eventually how defects affect the bending one-dimensional equation.

### 4.3 Boundary conditions for the bilayer

Boundary conditions concern the top and the bottom interfaces between the surrounding fluid and the two layers, as well as the interface between the two layers (*Z* = 0). In solid mechanics, the interface between layers is often considered as a line of discontinuity. More realistically, it is a thin zone of sharp variations for physical constants - in our case for growth coefficients or stiffness. Moreover, it may happen that the elasticity of the interface is not simply the mean-value of both elastic coefficients. As discussed above, this happens when the interface of a cell layer and the ECM is enriched in intracellular cytoskeletal filaments, or when a thin basal lamina separates two cell layers, both circumstances leading to a stiffer interface. If the interface is really smaller than each layer, we can consider it as a transition zone around *Z* = 0, then avoiding the application of boundary conditions to a system of 3 layers. A way to avoid writing boundary conditions is to represent the sample as a unique layer where the elastic parameters such as the Young modulus vary continuously. If the interface is very thin, one needs only to know its thickness relative to the thickness of the bilayer, the detail of the description is not really important. The fact that we use the so-called membrane hypothesis with cancellation of the stresses *σ*_13_ = *σ*_23_ = *σ*_33_ =0 ensure automatically the continuity of the normal and shear stress at the upper and lower boundaries and at the interface of the two layers (*Z* = 0). So only the continuity of the displacements is required at the interface. Choosing a “continuous approach” and aiming to transform the bilayer into a unique layer whose entire thickness will be *h*_*f*_ + *h*_*s*_, we assume that the Young modulus *E* and any growth number *g*_*α,β*_ can be written as:

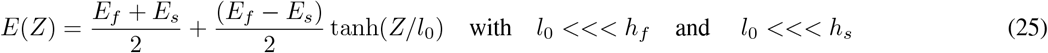

and

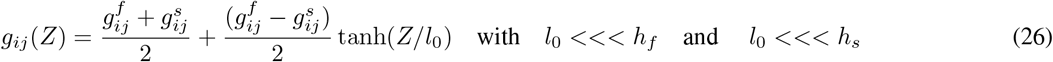

When a thin stiff membrane superposes to the two main layers of the sample at a position *Z*_*m*_ which can be on top *Z*_*m*_ = *h*_*f*_, or at the bottom *Z*_*m*_ = -*h*_*s*_, or at the surface of separation: *Z*_*m*_ = 0, as justified above, then we can slightly modify eq.(25) into

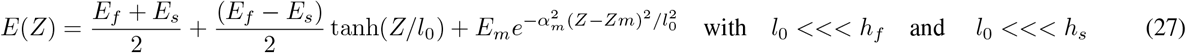

where the Young modulus of the membrane, *E*_*m*_, can be much larger than both *E*_*f*_ and *E*_*s*_. *α*_*m*_ is a numerical coefficient characterizing the thickness of the membrane. Another boundary layer may exist on top or bottom of the bilayer, in addition to the interfacial one. The formulation of the Young modulus, *E*(*Z*), must then also include this new contribution. Such a definition of *E*(*Z*) can be implemented into eq.(18) very easily to deliver the new FvK equations. We focus now on the neutral surface position and on the bending coefficients when either the elastic coefficients or the thicknesses present a sharp but small in size variation along the sample.

## 5 Position of the neutral surface *Z*_*n*_ for uniaxial loading

We can guess from the definition of the coefficients (eq.(24)) of the bending FvK equation (eq.(23)), that the position of the neutral surface is crucial for the shape of the sample when growing and buckling. We consider first the position of the neutral surface for layers of constant thickness and constant Young modulus and we introduce a linear perturbation dependent on the *x* variable. If the linear approximation is not valid, a numerical solution is always possible as in sect.(7).

### 5.1 Position of the neutral surface for a bilayer without structural defects

In the case of an ideal bilayer with no structural defects, 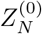 is determined implicitly via eq.(13) with a Young modulus *E*(*Z*) defined by: *E*(*Z*)= *E*_*f*_ if 0 ≤ *Z* ≤ *h*_*f*_ and *E*(*Z*)= *E*_*s*_ if -*h*_*s*_ ≤ *Z* ≤ 0. D is deduced from eq.(18) and it reads:

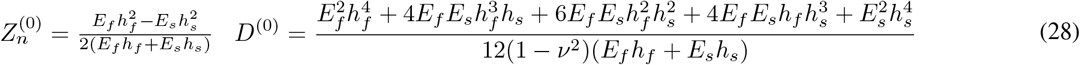

where the superscript “^(0)^” for *Z*_*n*_ and *D* reminds us that these formulas are restricted to ideal cases. One can notice that if the two layers have the same elastic coefficient, one recovers the standard bending stiffness: *D* = *E*(*h*_*s*_ + *h*_*f*_)^3^/(12(1 − ν^2^)) and the neutral surface is located at the middle of the layer of thickness (*h*_*f*_ + *h*_*s*_)/2. We can convince ourselves that the neutral layer is located inside the bilayer since 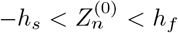, which is a necessary condition. These two quantities 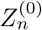 and D^(0)^ play a deep role in the bending equation, first equation of the FvK set of equation eq.(21) and for the definition given by eq.(18). It is why we consider now the departure from the ideal situation:

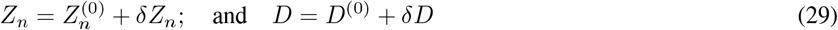

Hereafter, several causes are investigated at linear order in the perturbation amplitude.

### 5.2 Position of the neutral surface for diffuse or stiff interfaces

We allow a diffuse interface according to the representation given by eq.(25). The thickness of the interface *l*_0_ is small compared to *h*_*f*_ or *h*_*s*_. Although an exact calculation of *Z*_*n*_ is doable, it involves unusual analytical functions so we give here only an asymptotic formula for the result up to a correction of order 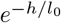 :

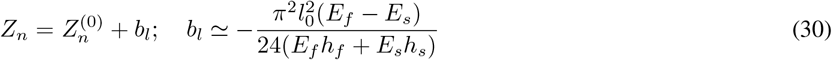

This is a tiny effect being of order 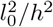. In the presence of a stiff interface with *E*_*I*_ larger than *E*_*f*_ or *E*_*s*_, the correction to *Z*_*n*_ is more important:

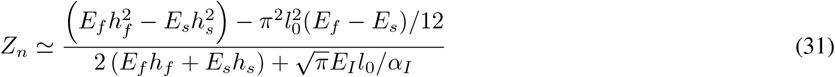

where the symbol ≃ indicates also a correction of order 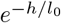, *h* being either *h*_*f*_ or *h*_*s*_. In the presence of an additional stiff layer, on top or on the bottom, with a different stiffness *E*_*b*_, we derive in the limit of vanishing 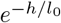, the following value for the neutral surface position:

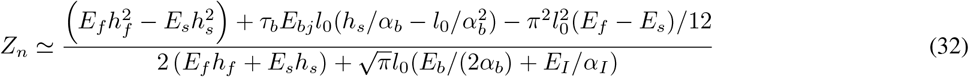

with *τ*_*b*_ =1 when the stiff layer is on top, and *τ*_*b*_ = − 1 when the stiff layer is at the bottom. Knowing that the interface stiffness *E*_*I*_ or *E*_*b*_ may be an order of magnitude larger than *E*_*f*_ or *E*_*s*_, this correction may modify significantly the values of the *D* coefficient in the FvK equations. The addition of thin layers, as described here, to the initial bilayered system will change the position of the neutral surface and the resulting *D* coefficient, but it will not change the structure of the FvK equations.

### 5.3 Localized defects of the thickness

Let us consider now a weak and localized variation of the thickness of one of the two layers. The neutral surface will then be distorted. We derive its new position by a simple linear expansion of *Z*_*n*_. This induces an equivalent expansion on the parameter *D* which multiplies the bending term in the equation eq.(23). Assuming that the thickness varies as

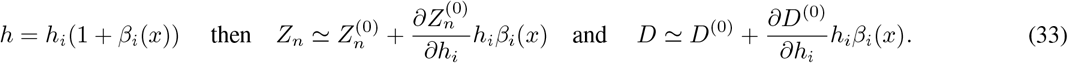

where D^(0)^ and 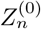 are given by eq.(28). and ≃ means an expansion restricted to first order. It reads

− On the top layer, we get:

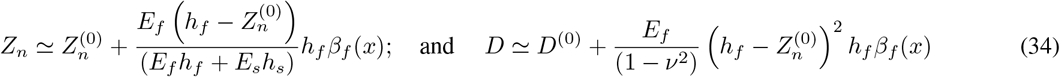

− On the lower layer:

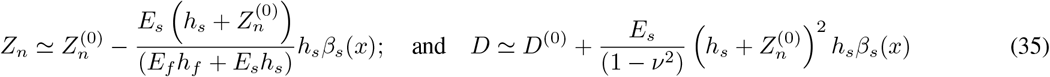

It is to be noted that a dip in either the upper or the lower layer means a negative value for the coefficients *β*_*i*_(*x*) so, as expected, a local decrease of the stiffness of the sample.

### 5.4 Localized inhomogeneity of the Young modulus

We now consider a localized variation of the Young modulus, which we treat through the expansion:

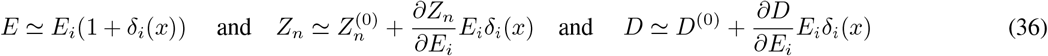

Leading to:

**–** Inhomogeneity in the top layer:

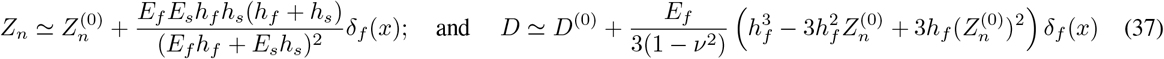

– Inhomogeneity in the bottom layer:

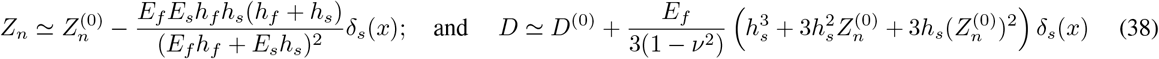

## 6 Results for uni-axial folding

As mentioned above, the uniaxial case simplifies a lot the analytical analysis but reduces the diversity of buckling patterns that can be accounted for compared to the 2D case. Nevertheless, the 1D approximation is still of great experimental relevance for some experimental systems. In the case of the experiments of fig.(2), for example, the tissue and the genetic perturbation can be assumed to be spatially invariant along the y-coordinate, making it essentially a 1D problem. In this context, the present section provides an in-depth analysis of bending contributions of the bilayer in the uniaxial case, with and without structural defects. We first analyze the bending equation of the uni-axial folding analytically. We then address the problem numerically by selecting possible 1D examples using the software of resolution Mathematica. Finally, simulations from the FEM (Finite-element method) software COMSOL Multi-Physics are presented.

### 6.1 Source of bending in bilayers

We will demonstrate in the following that the source of bending for growing bilayers are numerous and diverse and may have different biological origins. We analyze some cases here as examples. First, we define dimensionless parameters which will govern our FvK equations for a bilayer made of two layers with constant elastic coefficients.

#### 6.1.1 Dimensionless bending equation

Using as length unit the length *L* of the sample, eq.(23) reads:

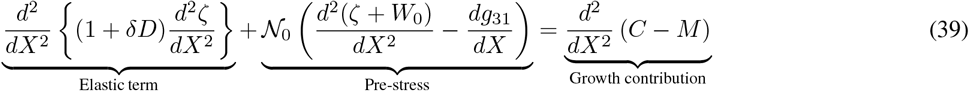

where we keep the same notation for *ζ*(*X*) which becomes *ζ*(*x*)/*L, X* = *x*/*L*, 𝒩_0_ = N_0_L^2^/D^(0)^, *δD* = (*D* - *D*^(0)^)/*D*^(0)^,*M* = *M*_11_*L*/*D*^(0)^,*C* = *C*_11_*L*/*D*^(0)^ are dimensionless quantities and *W*_0_(*X*)= *w*_0_(*x*)/*L* represents the initial position of the curved sample. External vertical loading is discarded. It is worth noting however that the equivalent of a loading can arise from the growth contribution (right-hand term of eq.(39)) owing to the fact that it involves spatial second derivatives, like the pre-stress term. For such a growth-induced loading to arise, growth of the two layers must differ to get a non-vanishing *M* value, and at least one of the terms (C or M) must vary spatially. This growth induced loading accords with previous investigations that demonstrated the need for differential growth of apposed layers to induce buckling of brain cortical folds for example [40–43]. Even with homogeneous growth, the right hand term can lead to a non-vanishing contribution, for example in the case of a structural defect that will change *Z*_*n*_, and induce high spatial frequency components in *C* or *M*. Nevertheless, it is difficult to discuss a priori the *ζ* profile which is added to *W*_0_ as a result of the growth term, since it depends also on the boundary conditions. We can intuitively predict that inhomogeneous growth is responsible for a “fictitious pressure” or on the contrary to a tension added to the sample. The simplest case may be the one of a homogeneous bilayer that we consider first.

#### 6.1.2 Bilayer without structural defects

This case corresponds to sect.(5.1). For simplification, we do not consider the case of a sharp interface and the bilayer is made of two perfect layers of constant thickness and Young modulus. The growth process occurs mostly in one layer. Assuming first 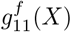 in the upper layer and 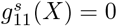 in the lower layer, we have for 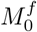 and 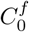 according to eq.(24):

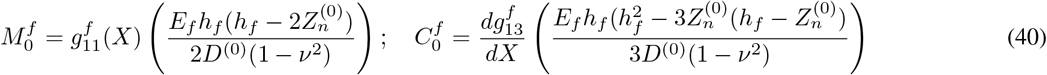

where *Z*^(0)^ and *D*^(0)^ are given by eq.(28). It is easy to check that both parameters *M*_0_ and *C*_0_ have the same sign as the growth elements 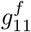 and 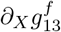. If growth occurs in the lower layer and not in the upper layer, it reads

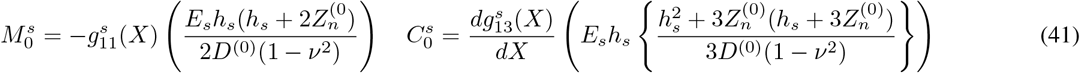

Here again, 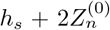 is a positive quantity and the sign of 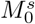 is opposite to the sign of 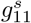 while the sign of 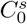 is automatically given by the growth coefficient derivative. Without pre-stress it is possible to observe a buckling of an initially flat plate because of a differential and inhomogeneous growth process. There exists a competition of the origin of this buckling between *g*_11_ or *g*_13_, or between *M*_0_ and *C*_0_. Differences in growth of the two layers induces a symmetry breaking between up and down. It is non trivial to compute the sign of the growth term -whether it contributes to a vertical pressure or tension-owing to the fact that it stems from the competition of two terms (*M*_0_ and *C*_0_) through their second derivatives. Notice however, if the growth is not *x* dependent, these terms will disappear from the buckling equation, on the contrary they will become more efficient if they strongly depend on *x*. Localized defects will increase the efficiency of the buckling as shown hereafter.

#### 6.1.3 Analysis of structural defect in growing bilayers

##### a) Localized stiffness variation in one layer

We first consider that the stiffness can be locally modified as discussed in fig.(2), inducing a small change in the position of the neutral surface. We focus on the linear variation of *M*^*f*^ or *M*^*s*^, and *C*^*f*^ or *C*^*s*^, simply deduced from eq.(40,41). The modification of these quantities are simply deduced taking into account the variation of *δZ*_*n*_ reported by eq.(37,38) in sect.(5.4), which reads:

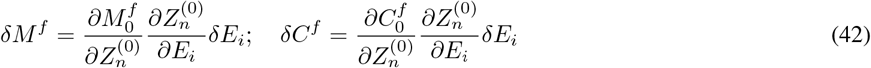

where *E*_*i*_ is the Young modulus of the layer affected by the defect: *E*_*f*_ or *E*_*s*_ and *δE*_*i*_ is represented by *E*_*i*_*δ*_*i*_(*X*), in a similar way to eq.(37,38). Similar results apply for *δM*^*s*^ which corresponds to growth in the lower layer and for *δC*^*s*^. We give here only the results but we define first the following positive quantities:

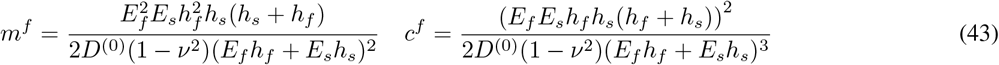

**–** When the growth occurs in the upper layer, which also exhibits a stiffness defect, then

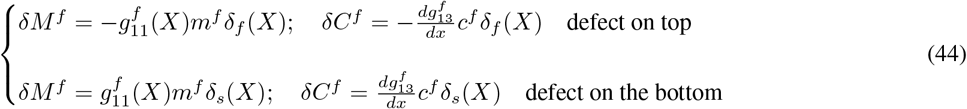

**–** When the growth occurs in the substrate, perturbation occurs now on 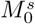 and 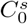 and one needs to consider again a defect of the stiffness either on top or on the lower layer.

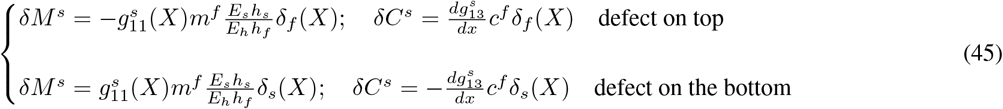

##### b) Localized variation of the thickness of one layer

A defect in the thickness affects the position of the neutral surface but also, when the thickness defect occurs in the same layer as the growth, the averaged value of the coefficients of eq.(24) related to the first bending FvK equation, eq.(23). We give here only the schema when a dip appears in the lower layer, the growth occurring either on the top or the lower layer. A similar analysis gives

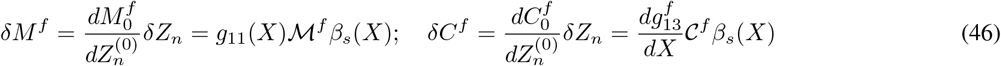

where as in sect.(5.3), *β*_*s*_(*X*) describes the shape of a notch and *ℳ*^*f*^ and *𝒞*^*f*^ are dimensionless positive quantities.

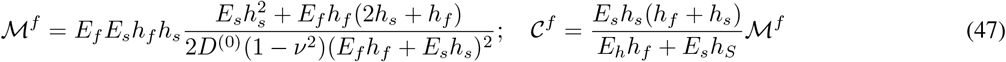

We examine now the case where growth occurs in the bottom layer where also the defect is localized: .

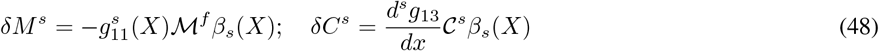

with

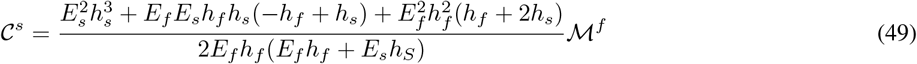

##### c) Shape of defects

Localized defects can be positioned in any place of the sample. From eq.(39), we have noticed that they may induce an equivalent forcing on the sample if they contribute to strong variation of the second derivatives of the coefficients *δM* and/or *δC*. Choosing a sharp defect in the center of the sample, a good representation may be 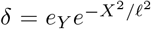where *e*_*Y*_ is the relative amplitude of the stiffness defect which can be positive or negative, 2𝓁 depicts qualitatively its width. The second derivative is *∂*_*XX*_*δ* is of order −2*e*_*Y*_ /𝓁^2^ which for a defect of width 0.05 corresponds to an amplitude of 200*e*_*Y*_ at the center. It is to be noted that the sign is opposite to *e*_*Y*_. So the sign of the second derivative of *δM* or *δC* can be easily found from paragraph (a). The convention for thickness defect is the following*:h*_*i*_*β*_*i*_(*X*) where *β*_*i*_(*X*) is negative for a notch and positive for a protrusion. Here again, *∂*_*xx*_*β* will be opposite to *β*. In any case, it remains that the prediction of the global sign of the second member of eq.(39) is hard to predict due to the competition between *g*_11_ and *g*_13_. Let us remember that a positive sign of a localized contribution will be equivalent to a localized forcing which will mimic the experiment [39] shown in fig.(3).

**Fig. 3.**
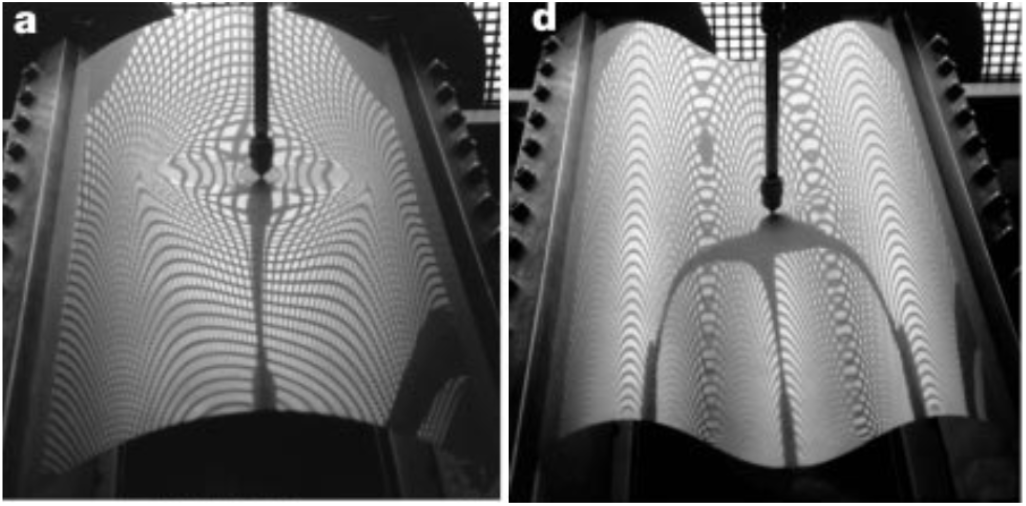
Cylindrical mylar shell under a localized forcing. Originally the plate has a length of *λ* = 35 cm and a width *L* = 17.5 cm. The Young modulus of the mylar plate is 3.810^9^ Pa.s and the Poisson ratio is *ν* = 0.4. The initial bending is realized by imposing the boundary conditions. On the left, one notices that for a low value of the imposed deviation *Z* = 4 mm, the pattern of sinking is quite elliptical while it becomes cylindrical for *Z* = 15 mm representing a folding in the opposite direction of the initial one. Image extracted from [39].

### 6.2 Numerical investigations of the FvK bilayer for a plate or a shell

In this section, we aim to demonstrate the role of defects on plates and slightly curved shell portions under pre-stress. To simplify we focus first on a homogeneous growth process: it means a process independent of *X* and identical in both layers or equivalently a non growing sample. Then we will give numerical examples to illustrate the analytical results of the previous section when differential growth occurs contingently to defects.

#### 6.2.1 Decrease of thickness on a non-growing pre-stressed plate or shell

In this section, we will evaluate the role of a sharp variation on the *D* coefficient of a pre-stressed sample initially flat or curved. For numerical purpose the defect has the shape of a penetrating peak and is represented by a Gaussian so 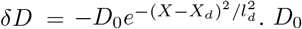. *D*_0_ is a positive coefficient which can be evaluated with either eq.(34) or eq.(35), *l*_*d*_ is an estimation of the scale of the defect compared to the scale of the sample *L* and *X*_*d*_ is its relative position in the sample. For a plate, a curvature is initially induced by the pre-stress and applying a slope at both ends. In fig.(4a) the imposed slope is *∂*_*X*_*ζ* ± 0.03 at *X* ± 0.5 giving a deflection below the horizontal (black line). This deviation increases with the defect since it weakens the stiffness more and more with the amplitude *D*_0_ = 0.2 (blue curve), *D*_0_ = 0.3 amplitude (red curve). Even for a sharp-pointed defect, with *l*_*d*_ = 1/10, the deviation *ζ* remains small and not very selective. In addition, a displacement on the position of the defect does not affect *ζ* too much (magenta curve). For a shell which is initially curved in the positive *Z* direction with clamped conditions at both ends, as shown in fig.(4b,c), the deviation *ζ* is amplified by the initial curvature and the pre-stress. Here again, any deviation on the right of the defect weakens its role on *ζ* since we cannot distinguish the deviation with lateral defect (magenta line) from the black line. When the whole profile *Ω* = *ζ* + *w*_0_(*X*) is considered, we clearly observe an increase of the total curvature when the defect is centered at the maximum of deflection.

**Fig. 4.**
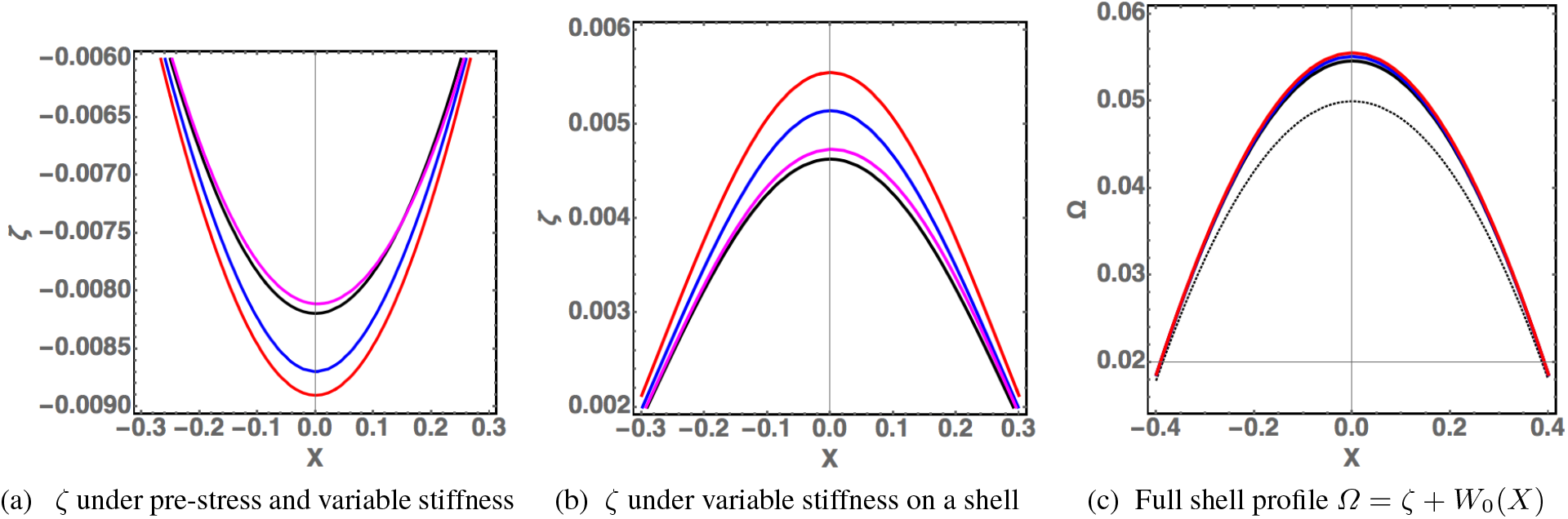
Buckling deviation due to a variable stiffness coefficient *D*(*X*), in a *pre-stressed non-growing* plate or shell. In **(a)**, the profile *ζ* of an initial elastic plate under pre-stress *N*_0_ = 4 is plotted as a function of the *X* coordinate. The *X*-domain is reduced to the interval *{*− 0.3, 0.3*}* for clarity. At both ends,*X* ± 0.5, *ζ* = 0 and 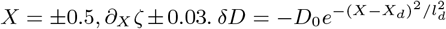 with *D*_0_ = 0 for the black line and *D*_0_ = 0.3 for the blue, red, magenta colored lines. Except for the magenta line, the dip is centered on *X* = 0, otherwise, it is located at *X*_*d*_ = 0.3. The blue and magenta curves correspond to a narrow dip with *l*_*d*_ = 0.1, the red curve to a larger one with *l*_*d*_ = 0.3. Notice that a dip located far from the center has no effect and the deflection remains weak in any case (scale of the vertical line). In **(b)**, *ζ* for a pre-stressed shell of equation *W*_0_(*X*) = (1/4 −*x*^2^)/5 of negative curvature and clamped boundary conditions :*X* ± 1/2,*ζ* = *∂*_*X*_ *ζ* = 0. In **(c)** the full profile *Ω* = *ζ* + *W*_0_ with a dotted black line for *W*_0_. The same code of colors is applied in the three panels **(a, b, c)**.

#### 6.2.2 Thickness defects on growing plates and shells

The thickness variation on *δD* gives a very intuitive result. A much less intuitive result concerns the sign of 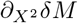 and 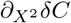 of eq.(39) since the results are dependent of the second derivative of these growth coefficients. These quantities have been studied in detail in sect.(6.1.3). We must notice that they have an opposite sign a priori and in practical situations, we will have little information except by comparison with genetically modified tissues. So we will join these two contributions into a unique term 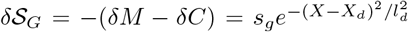 in the right-hand-side of eq.(39). Intuitively, as soon as this term is differentiated 2 times, it behaves as either a pressure if it is positive or a tension in the opposite case while the initial curvature of the concave shell behaves like a pressure and a convex one like a tension. Knowing that a groove in the substrate is represented by a negative profile, the second derivative is then positive for *X* = *X*_*d*_ being given by 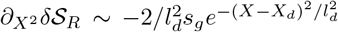 Illustration is given in fig.(5) for a pre-stress plate with defect localized at *X* =0 with two different length-scales: *l*_*d*_ = 0.1 and *l*_*d*_ = 0.3 and different amplitudes for *s*_*g*_ = ± 0.5 and ±1. A defect put on the right is also shown with a noticeable distortion in panel (c) of fig.(5). In fig.(6), the same set of perturbations act on a shell either concave (panel(a)) or convex (panel(b)) and the deviation *ζ* is plotted with boundary conditions *ζ* = *∂*_*X*_*ζ* = 0 for *X* ± 0.5. For the concave case (a), the deviation due to the defect is decreased for *s*_*g*_ > 0 and increased otherwise. The opposite result is obtained for in the convex case panel (b).The distortion is shown in panel (c) only for the concave case. Once the deviation is superposed on the initial shell, depending on the respective sign of the deviation versus the shell geometry, it is possible to observe an inversion of the curvature. This is demonstrated in the three panels of fig.(7).

**Fig. 5.**
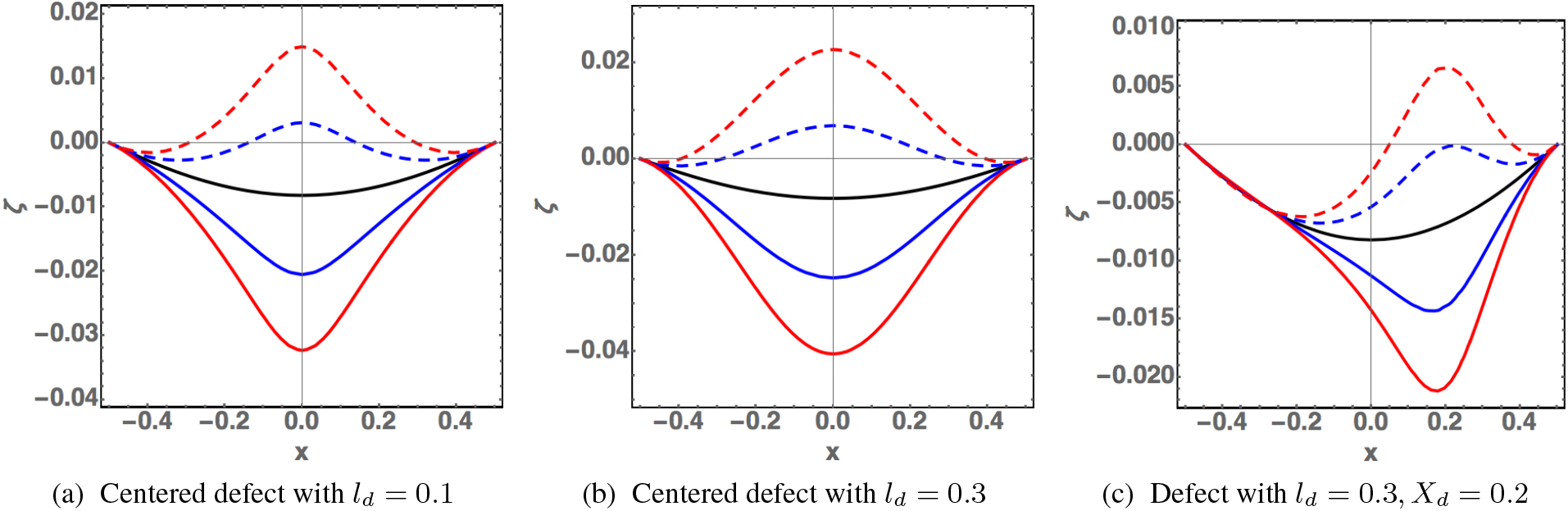
Buckling deviation *ζ* in a pre-stressed *growing* plate. The prescribed boundary conditions are: *ζ* = 0 and *∂*_*X*_ *ζ* = *±*0.03 at *X ±* 0.5, *N*_0_ = 4 and the black line is the initial shape of the plate under pre-stress. The defect is as follows: for *D*, 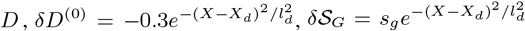, with *s*_*g*_ = 0.5 (resp. −0.5) represented by a continuous blue line (resp. a dot-dashed blue line) and *s*_*g*_ = 1 (resp. *s*_*g*_ = − 1) represented by a continuous red line (resp. dotted-dashed red line) for the 3 panels. In **(a)**, *X*_*d*_ = 0 and *l*_*d*_ = 0.1. In **(b)**,*X*_*d*_ = 0 and *l*_*d*_ = 0.3. In **(c)** the same as in **(a)** for an eccentric defect positioned at *X*_*d*_ = 0.2. Notice the strong asymmetry of all curves in panel (c).

**Fig. 6.**
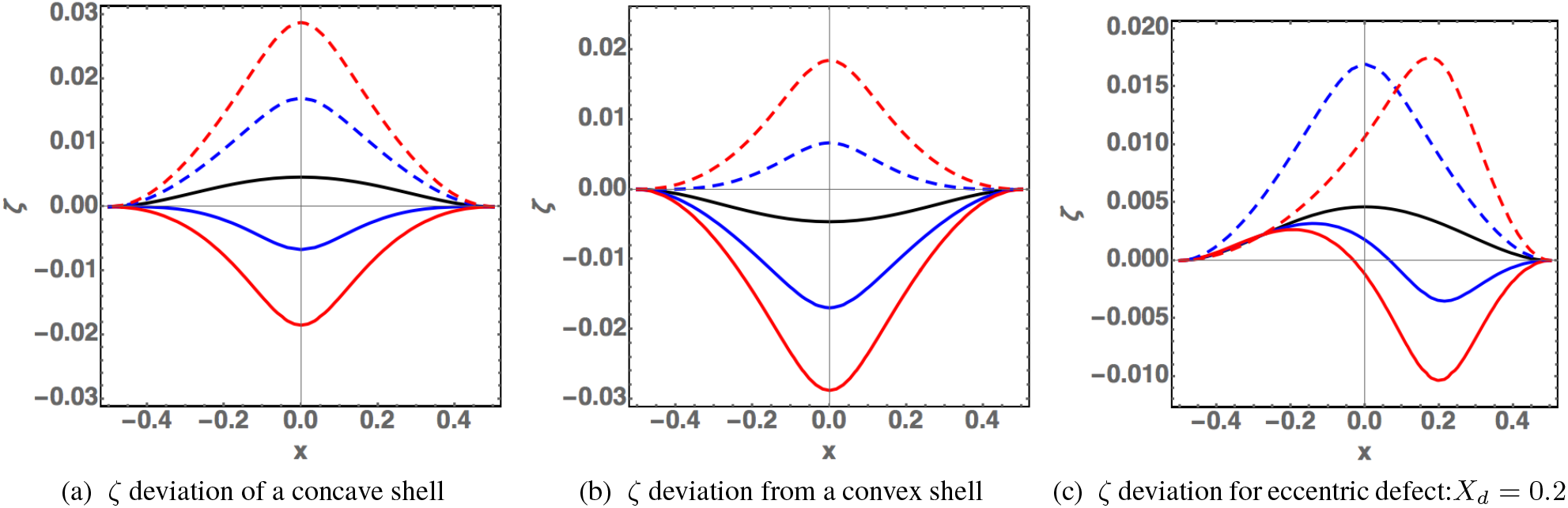
Buckling deviation *ζ* for a pre-stressed (*N*_0_ = 4) shell. The same definition and color codes apply as in fig.(5): black no defect, blue continuous *s*_*g*_ =0.5,blue discontinuous *s*_*g*_ = − 0.5, red continuous curve *s*_*g*_ = 1 and *l*_*d*_ = 0.1, red discontinuous *s*_*g*_ =_1_ for the 3 panels. Clamped boundary conditions are applied. In **(a)**, the shell is initially concave and *l*_*d*_ = 0.1. In **(b)** the shell is initially convex. In **(c)**, the defect is localized on the right *x*_*d*_ = 0.2 and the shell is concave. *W*_0_(*X*) = (1/4 − *X*^2^)/5., for concave shape and *W*_0_(*X*) = (*X*^2^ − 1/4)/5 for convex shape

**Fig. 7.**
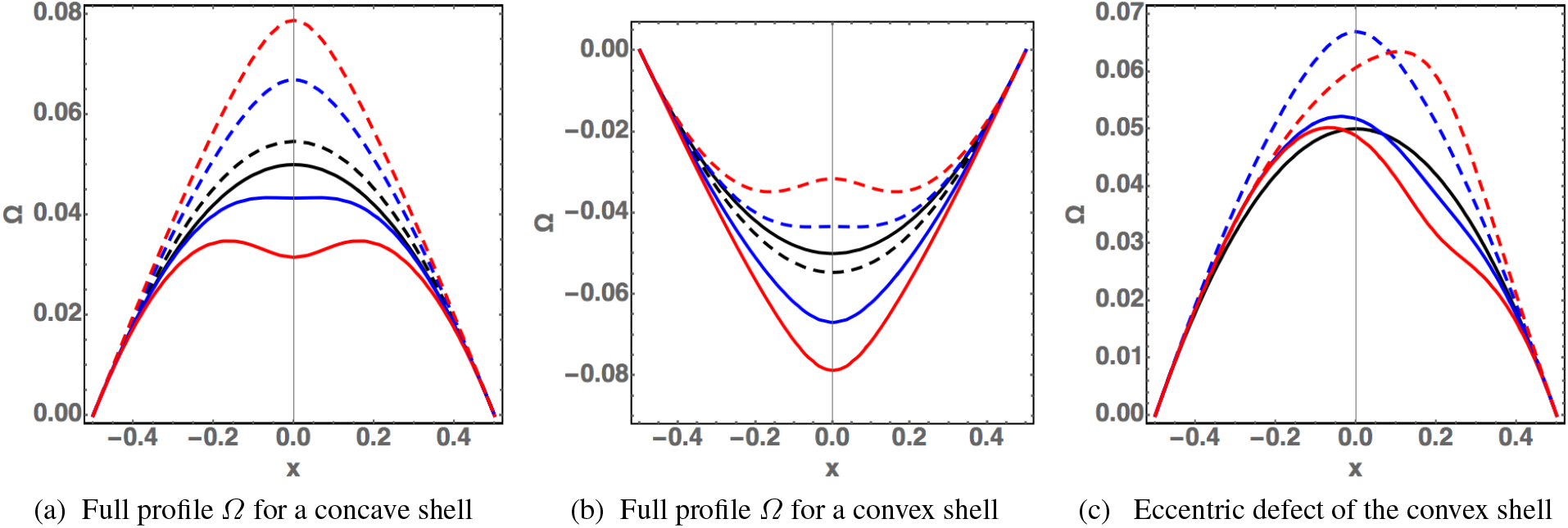
In **(a**,**b**,**c)**The shell profile *Ω*(*X*) = *W*_0_(*X*)+ *ζ*(*X*). The colored lines correspond to the buckling conditions of fig.(6). Notice the possibility to observe an inversion of curvature in panel (**a**) and (**b**) for a suitable size of *s*_*g*_ with respect to the curvature of the shell.

### 6.3 Confrontation of the uniaxial FvK bilayer model with experiments

In section (2), we presented experimental evidence that a local change in the mechanical properties of the ECM can induce major morphological changes in the wing imaginal disc epithelium. Here, we confront our bilayer FvK formalism in the uniaxial geometry to experimental profiles extracted from the images of fig.(2c,e).

We first focus on the tissue presented in fig.(2c), that we rescale in the unit of the lateral size of the imaginal disc (fig.(8a)). The perturbation of the ECM was induced before the naturally occurring curvature could develop in the tissue. We therefore assume no initial curvature: a horizontal plate only under pre-stress. We fix the slope conditions at the border, making the hypothesis that the tissue periphery is not affected by the perturbation. This hypothesis is justified by the fact that the defect is well localized at the center of the tissue. The fit of parameters is achieved by a limited number of trials. Fitted parameters converge to the following values: a pre-stress of order 9 in unit of *D* and a parameter *D*_0_ = 0.3; a defect characterized by an amplitude *s*_*g*_ = 4, position *x*_*d*_ = 0.1 and width *l*_*d*_ = 0.25, when the defect is modeled by the relation 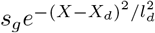, see sect.(6.2.2). Finally in fig.(8b), the shape of the disc at the boundaries and the developmental timing of the perturbation, suggest that the tissue was already curved at the onset of the perturbation. We then used the shell treatment with a best guess for the unperturbed shape *w*_0_(*X*)= 0.7(1/4 − *X*^2^ + 4(1/16 − *X*^4^)), *N*_0_ has little effect and is put to zero, which suggests that in old epithelia, the outer stresses have relaxed which is not the case for young epithelia. The amplitude of the defect has increased: it is now *s*_*G*_ = 10; localization is *X*_*d*_ = − 0.05, and width *l*_*d*_ = 0.1. Thus, the bilayered FvK theory can account for the mechanics at play in the growing wing imaginal disc in the presence of a localized defect in the ECM, both when the tissue is initially flat and curved.

**Fig. 8.**
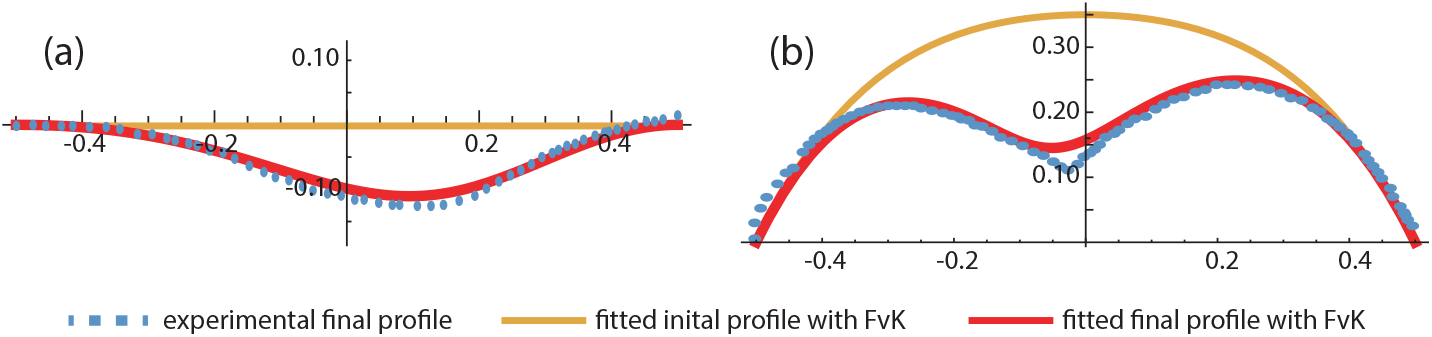
Fitting experimental wing disc profiles with the uniaxial FvK bilayer theory. **(a)**: Theoretical fit of an early wing imaginal disc, for which the initial profile before perturbation was flat (same as fig.(2c)). **(b)**: Theoretical fit of an older wing imaginal disc, for which the initial profile before perturbation was curved (same as fig.(2e).

## 7 Numerical Simulations with Finite-Element Method

We now consider the problem of the bilayer bending in the linear Hookean elasticity approach using Comsol Multiphysics which computes the deformations in the real geometry of the bilayer, two different connected layers, without averaging the elastic properties. The bilayer we study in the following is made of a thin (*h*_*f*_ = 10μm) and relatively soft (*E*_*f*_ = 50 kPa) upper layer and a thicker (*h*_*s*_ = 12μm) and stiffer (*E*_*s*_ = 75 kPa) bottom layer. The length of the bilayer is *L* = 500 μm. The layers are submitted to growth and pre-stress, and the boundaries are either free or constrained. We investigate a defect, as a local removal of the basal layer in the middle of the bilayer with an extension of 10 μm and a thickness *h*_*d*_ ≤ *h*_*s*_, and as a local change of stiffness and growth in the bilayer. The growth is anisotropic and is introduced in a tensorial form: *G*_*ij*_ = 1 + *g δ*_1*i*_*δ*_1*j*_, with *g* ∼ 0 − 0.1. The growth may be introduced in the top layer only as in sect.(7.2), the bottom layer only in sect.(7.4), or in both layers as in sect.(7.3). The pre-stress consists in an anisotropic compression or tension, so that 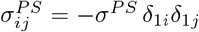 where *σ*^*PS*^ ∼ 0 − 500 Pa.

We also study the case of the shell on the further deviation of the sample. We set an initial curvature to the shell: *W*_0_ = *H* (*L*/2)^−2^((*L*/2)^2^ *X*^2^) where *H* is the displacement at *X* = 0. In order to compare the different results, we plot the profile of the interface between the upper and lower layer (the red line in fig.(9e).

**Fig. 9.**
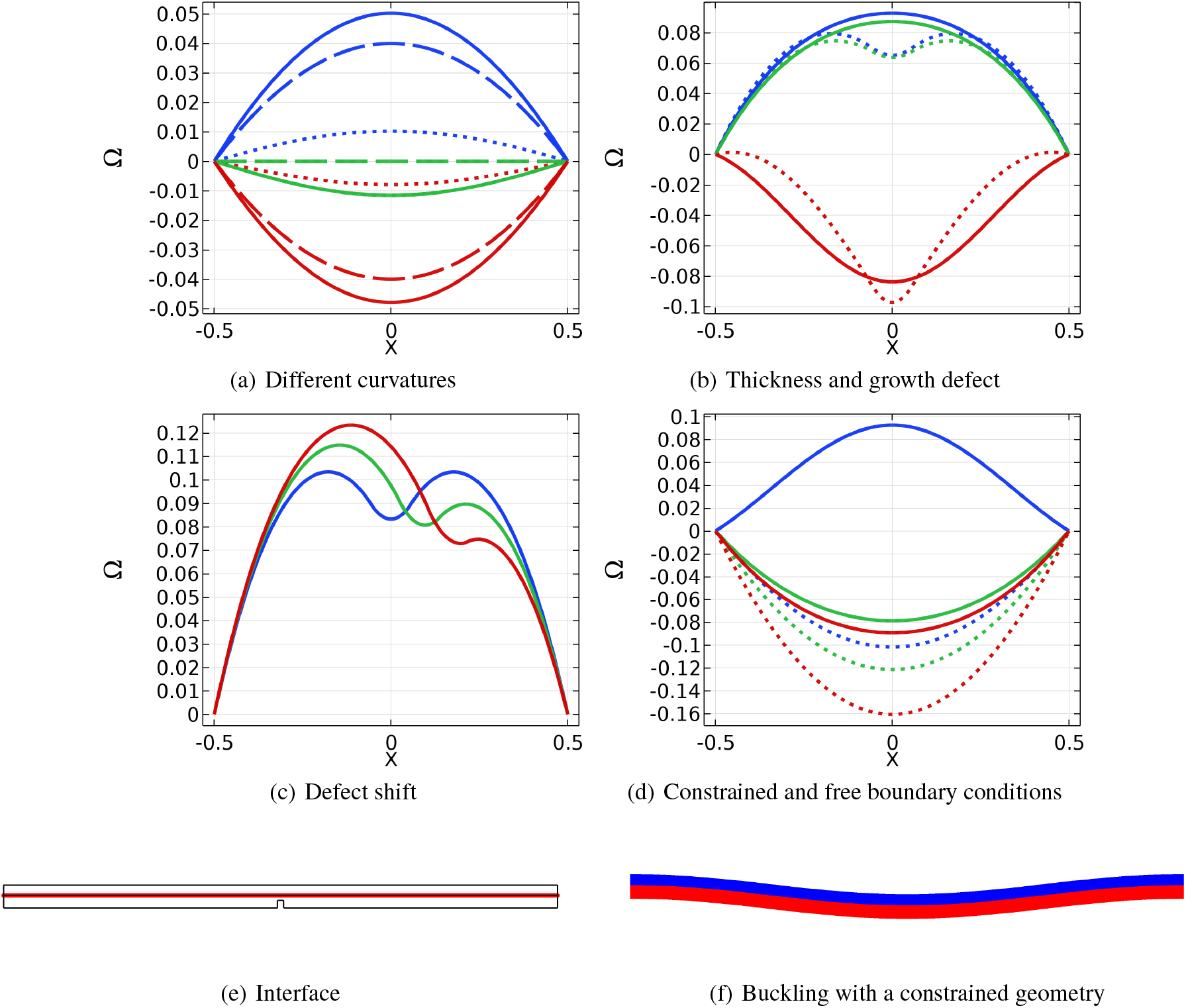
**(a):** Profile of the interface between the two layers for different curvatures and a pre-stress *σ*^*PS*^ = 200 Pa, with no defect in the lower layer. The continuous, dashed, and dotted lines correspond respectively to *W*_*0*_ + ζ, *W*_*0*_ and ζ. The red, green and blue lines correspond respectively to *H* = 20, 0, 20 μm. **(b):** Profile of the interface between the two layers for a case with a defect of relative thickness *h*_*d*_*/h*_*s*_ = 0.85 with *h*_*d*_ the defect thickness and *h*_*s*_ the bottom layer thickness. The growth parameter is *g* = 0.05. The green, blue, and red curves correspond respectively to cases with an initial zero, concave and convex curvatures *H* = 0, 20, 20. **(c):** Different shifts for the defect, which is at *X* = 0, 0.08, 0.16 respectively for the blue, green, and red line. The amplitude of the deformation depends on the localization of the defect. The growth parameter takes the value *g* = 0.1. **(d):** Comparison between constrained (dashed lines) and free (continuous lines) boundary conditions for zero, convex and concave initial curvatures (*H* = 0, 20, 20 for the green, red and blue lines) and a growing bottom layer (*g* = 0.03). **(e):** Interface between the lower and upper layer in red. **(f):** Buckling for a constrained bilayer with a growing bottom layer and an initial concave curvature.

Finally, we fit FEM simulations to the experiment described in sect.(2). The result is compared to the one obtained from the FvK calculation sect.(6.3).

### 7.1 Pre-stress on an initially curved bilayer

In fig.(9a) we investigate different initial shell shapes with different curvatures characterized by *H* = −20 μm, 0 μm, 20 μm with the same pre-stress *σ*^*PS*^ = 200 Pa in the bilayer. For a plate, with no initial curvature, a bending toward the stiffer and thicker basal side is obtained while for a convex or a concave shell, the initial curvature of the shell is reinforced in absolute value. This is consistent with the previous FvK model for shells. In fact the bending deviation follows the initial shape of the shell.

### 7.2 Thickness and growth defect on growing plates and shells

In biological systems, the two layers in epithelia are not independent. For instance, the basal membrane regulates the proliferation and the differentiation in the epithelium. In general, the basal layer structures the upper layer of the epithelium, and a defect in the lower layer can have consequences on the proliferation and metabolism of the upper layer. Therefore, we now investigate the shape change at the level of a defect which alters the thickness of the basal layer and consequently the growth of the upper growing layer. Figure (9b) displays the results when the basal layer is removed in proportion of 85% (dotted lines) in thickness and the upper layer is not growing at the defect level, as well as the case with no defect (continuous lines). We observe a buckling towards the basal membrane, in the neighborhood of the defect.

### 7.3 Place of a defect

Defects can appear anywhere along a bilayer. In fig.(9c) we impose defects on a planar bilayer, located at *X* = 0, 0.08, 0.16 for the blue, green and red lines respectively. Both layers grow to the same extent *g* = 0.1. The defect consists in removing a part of the bottom layer. Interestingly, with the same size of the defect, the amplitude of the deformation is different, and the shift of the defect is not only a shift in the deformation.

### 7.4 Dependence on boundary conditions

The comparison between confined and free geometries is a recurrent topic in cellular growth studies [44,45] from multiple perspectives, such as the folding of epithelial sheets [46]. The boundary conditions of bilayer plates or shells are also necessary for the analytical and numerical study of the system. But unfortunately, they are not always perfectly known. When the set-up is symmetric, one can use those symmetries to constrain the problem, and the system has free-boundary conditions. Sometimes the experiment is such that the system is clamped, or loses its symmetry. Confined systems are strongly dependent on boundary conditions.

In fig.(9d) we compare the buckling in a constrained and free geometry. The buckling is very different, and its direction (up or down) even changes when starting with a concave geometry. It may lead to strong inaccuracies in FEM simulations of rectangular geometry and semi-analytical treatments become necessary, see reference [47]. Notice that in the previous sections, sect.(6.2.1,6.2.2), boundary conditions are always fixed symmetrically for plates and clamped boundary conditions for shell deviations. This point is important for comparing our numerical results to the experiments as shown in the next section. There can be a few types of constrained geometry. In fig.(9d), it is obtained by imposing a zero displacement at two points on the lateral boundaries. In fig.(9f), the whole lateral boundaries are imposed a zero displacement. In the second case, the *Z*-displacement is close to having a zero derivative as boundary conditions.

### 7.5 Confrontation of the finite-element simulations with experiments

We use the finite element method to reproduce qualitatively the side to which the imaginal wing curves. We assume that the stress inside the cell layer is induced by growth, and is relaxed when a defect is introduced. Since in classical mechanics only the stress-free configuration and the configuration obtained after the process are to be taken into account, we make a multi-layer growth with and without defect. In our simulations, we observed that a 2-layer model could not reproduce the experimentally observed change of curvature upon ECM degradation. Only when an apical membrane is introduced in the model, can we reproduce the behavior - making it essentially a 3 layers model. Indeed, contrary to the resolution of the FvK equation where boundary conditions are imposed, we use free boundary conditions with the Finite Element Method. Therefore a third layer has to be added in order to constrain the system to bend towards the correct side.

More precisely, we simulated a thick (15 μm) cell layer of length *L* = 500 μm, growing between two non growing thin layers. This process creates a compressive stress in the two non growing layers. The top layer has a thickness of 5 μm and the lower layer has a thickness of 2 μm. The apical membrane may represent the apical cortex, and the apical adherens layer that dominates apical mechanics in epithelia [4], whereas the basal membrane represents the ECM. The growth deformation gradient is written as: *G*_*ij*_ =1 + *δ*_1*i*_*δ*_1*j*_*g*_11_. The boundary conditions are free.

We fit the experiment (fig.(11a,11b)) and we find a basal ECM stiffer (*E*_*b*_ = 100 kPa or *E*_*b*_ = 30 kPa) than the upper membrane and that the cell layer (*E* = 10 kPa). In the case of the early imaginal wing, *g*_11_ = 0.2 and *E*_*b*_ = 30 kPa, and for the old imaginal wing *g*_11_ = 0.25 and *E*_*b*_ = 100 kPa. We also introduce an asymmetry for the place of the defect at *X* = − 0.025 for the old epithelium and *X* = 0.15 for the early epithelium. The apparent growth is thus higher and the ECM stiffer for the old imaginal wing when compared to the early imaginal wing. This set-up results in a bending toward the upper layer in the case with no defect for the old imaginal wing and a slight bending towards the basal side for the early imaginal wing (fig.(10a,10c)). However, when a defect of length *l*_*d*_ = 100 μm is introduced in the form of a removal of the basal ECM, the curvature undergoes an inversion toward the basal side in in both cases (fig.(10b,10d)).

**Fig. 10.**
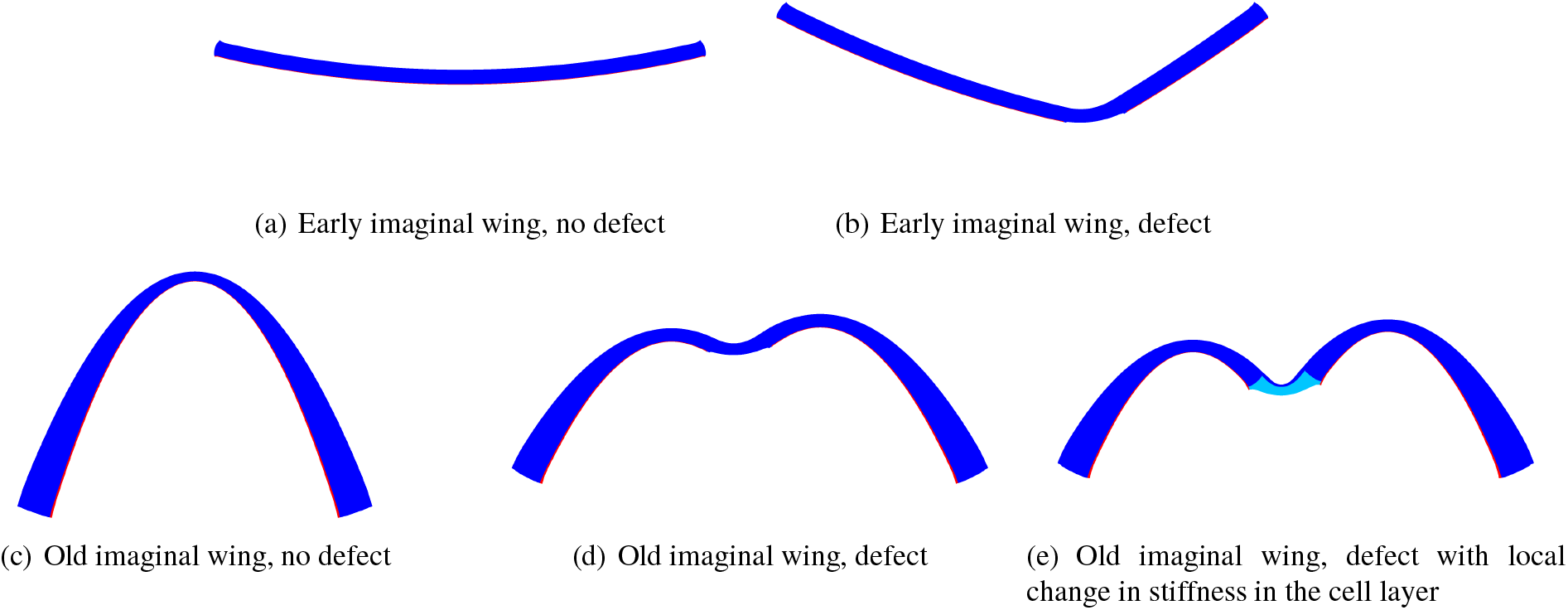
Cuts of the imaginal wing for the different FEM simulations with and without defect. The colors reflect the stiffness: red for the basal ECM, blue for the upper membrane and the cell layer, and cyan for the zone of the cell layer around the defect that softens in the refined model.

It seems that for the early imaginal disc this model is sufficient to properly reproduce the experimental profile (fig.(11a)). However, the agreement was not as good for the older initially curved imaginal disc (fig.(11b)).We then modified the FEM model with the same strategy as for the FvK model, by changing the stiffness at the level of the defect, making the cell layer softer. This refined model provided a better fit to the experiments (fig.(10e,11c)). The new values of the fit for the old imaginal wing are: *g*_11_ = 0.43 and *E* = 1.7 kPa in the cell layer at the defect level. The other parameters remain unchanged. Contrarily, changing the growth rate at the level of the defect does not improve the fit. We deduce that the contribution s_g_ of the defect in sect.(6.3) is mostly due to a change of the stiffness in the growing cell layer at the level of the defect, rather than a change of growth. This is consistent with the duration of Mmp2 expression during the experiment (18-24 h), see sect.(2). No pre-stress was introduced in the FEM simulations, since this pre-stress is assumed to be caused by growth, and the growth can be introduced explicitly with an order of magnitude larger than with FvK. For the same reason, growth generates the curvature, which is not introduced explicitly. To conclude, FEM simulations validate the FvK calculations.

**Fig. 11.**
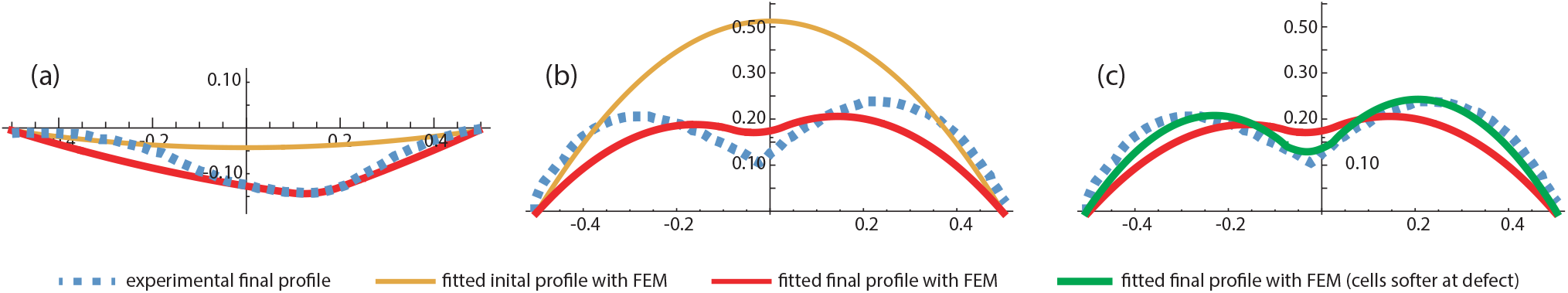
Fitting experimental wing disc profiles with the FEM model. **(a)**: Theoretical fit of an early wing imaginal disc, for which the initial profile before perturbation was flat (same as fig.(2c)). **(b)**: Theoretical fit of an older wing imaginal disc, for which the initial profile before perturbation was curved (same as fig.(2e). The stiffness of the cell layer is not altered by the defect. **(c)**: Comparison between the refined fit when the stiffness of the cell layer is altered by the defect (in green) and the case when it is not (red).

## 8 Discussion and conclusion

Motivated by experimental evidence that the ECM plays an important role in shaping epithelia in 3D, as shown in fig.(2), this work provides a mechanical description of growing epithelia in the Föppl-von Kármán framework. In the context of embryo-genesis and organo-genesis, previous theoretical studies have modeled epithelia using the vertex model [48–50] inspired from the physics of foams [51]. Indeed, epithelia are cellular pavings where the main unit has a polygonal shape in 2D or a shape of polyhedrons in 3D. Vertex models approximate cells as polygons, make hypotheses about the mechanics of individual cells (*e*.*g*.their growth, surface or line tension and stifness) and simulate the behavior of cell assemblies from these elementary rules. These simulations are based on energy minimisation, as in the present work. One limitation of cell-based models is the fact that simulating cell assemblies may become difficult at very large cell numbers, notably when the parameter-space must be explored. In addition, it is rather hard to extract macroscopic physical quantities such as Young modulus or surface tension from microscopic parameters describing the cells.

Most of the aforementioned theoretical works focused on the 2D mechanics of the apical surface of epithelia, with only few studies addressing the 3D aspects of tissues [52,53]. Epithelia are usually thin layers and the elasticity of slender elastic objects such as plates, tubes or membranes can be good candidates to describe their mechanical properties especially in the presence of bending. However, a single plate may be too simple a model when it comes to explaining the behavior of epithelia, which are intrinsically multilayered systems. In the present paper we have considered the multi-component structure of epithelia from the view point of elasticity. Local changes in elastic coefficients and differential growth are at the origin of stresses able to buckle the sample. Not only could our bilayered elastic model predict a broad range of morphogenetic behavior, it also yielded surface profiles that were in very good agreement with experimentally observed ones. Notably, it could account for the localized degradation of the ECM on two experimental observations that were limit cases: one initially flat and one initially curved wing imaginal discs (figs. (2,8)). The morphogenetic processes that we observed and modeled -fold formation upon ECM degradation-were artificially induced *via* an ectopic genetic expression of the metalloprotease Mmp2. These experiments, as such, fall in the realm of “synthetic morphogenesis”. Nevertheless, similar processes are naturally occurring in tissues. For example, deep folds that develop at late stage of the wing imaginal disc are thought to arise through such a local degradation of the ECM [23,24]. In this paper, we performed experimental observations of tissues exclusively after the action of the genetic perturbation. Ideally, one would need to image the tissues before and after the onset of the perturbation to completely disentangle the origin of the buckling. This is experimentally possible in *Drosophila* with chronic imaging [54] and potentially through prolonged imaging (*i*.*e* not just a few snapshots) with an appropriate method to reduce the phototoxicity associated with the light dose [55].

In the uniaxial geometry (with only one dimension for the sample), defects generate buckling distortions of the shape due to variation of thickness, stiffness but also of growth process. For a 1D bending process, the in-plane elastic equation is trivially solved and only the bending equation exhibits interesting degrees of distortions. It is not the case for 2D modes of deformations where spatial patterns happen on the outer periphery of the samples ([15,16]). One can then expect interesting patterns in these geometries.

## 9 Author Contributions

The two first authors contributed equally. Both LL and MBA conceived the project and wrote the manuscript.

## 10 Acknowledgements

The authors acknowledge the support of ANR (Agence Nationale de la Recherche) under the contracts MecaTiss (ANR-17-CE30-0007) and GuideFusion (ANR-18-CE13-028); Excellence Initiative of Aix-Marseille University - A*Midex (capostromex), a French Investissements d’Avenir programme. PQQ acknowledges the support of the China Scholarship Council (CSC), file *N*^0^ 201706100182.

## References

1. B. Li, Y.-P. Cao, and X.-Q. Feng, “Growth and surface folding of esophageal mucosa: a biomechanical model,” Journal of biomechanics, vol. 44, no. 1, pp. 182–188, 2011.

2. M. Ben Amar and F. Jia, “Anisotropic growth shapes intestinal tissues during embryogenesis,” Proceedings of the National Academy of Sciences, vol. 110, no. 26, pp. 10525–10530, 2013.

3. E. Hannezo, J. Prost, and J.-F. Joanny, “Instabilities of monolayered epithelia: shape and structure of villi and crypts,” Physical Review Letters, vol. 107, no. 7, p. 078104, 2011.

4. D. Pinheiro and Y. Bellaiche, “Mechanical force-driven adherens junction remodeling and epithelial dynamics,” Developmental cell, vol. 47, no. 1, pp. 3–19, 2018.

5. N. Khalilgharibi and Y. Mao, “To form and function: on the role of basement membrane mechanics in tissue development, homeostasis and disease,” Open Biology, vol. 11, no. 2, p. 200360, 2021.

6. P. Ciarletta and M. Ben Amar, “Papillary networks in the dermal–epidermal junction of skin: a biomechanical model,” Mechanics Research Communications, vol. 42, pp. 68–76, 2012.

7. J. Bateman, R. S. Reddy, H. Saito, and D. Van Vactor, “The receptor tyrosine phosphatase dlar and integrins organize actin filaments in the drosophila follicular epithelium,” Current Biology, vol. 11, no. 17, pp. 1317–1327, 2001.

8. I. Delon and N. H. Brown, “The integrin adhesion complex changes its composition and function during morphogenesis of an epithelium,” Journal of cell science, vol. 122, no. 23, pp. 4363–4374, 2009.

9. M. A. Biot, “Folding of a layered viscoelastic medium derived from an exact stability theory of a continuum under initial stress,” Quarterly of Applied Mathematics, vol. 17, no. 2, pp. 185–204, 1959.

10. M. Kücken and A. C. Newell, “A model for fingerprint formation,” EPL (Europhysics Letters), vol. 68, no. 1, p. 141, 2004.

11. M. Kücken and A. C. Newell, “Fingerprint formation,” Journal of theoretical biology, vol. 235, no. 1, pp. 71–83, 2005.

12. B. Audoly and A. Boudaoud, “Buckling of a stiff film bound to a compliant substrate—part iii:: Herringbone solutions at large buckling parameter,” Journal of the Mechanics and Physics of Solids, vol. 56, no. 7, pp. 2444–2458, 2008.

13. P. M. Reis, F. Corson, A. Boudaoud, and B. Roman, “Localization through surface folding in solid foams under compression,” Physical review letters, vol. 103, no. 4, p. 045501, 2009.

14. J. Yin, J. L. Yagüe, D. Eggenspieler, K. K. Gleason, and M. C. Boyce, “Deterministic order in surface micro-topologies through sequential wrinkling,” Advanced materials, vol. 24, no. 40, pp. 5441–5446, 2012.

15. J. Dervaux and M. Ben Amar, “Morphogenesis of growing soft tissues,” Physical review letters, vol. 101, no. 6, p. 068101, 2008.

16. J. Dervaux, P. Ciarletta, and M. Ben Amar, “Morphogenesis of thin hyperelastic plates: a constitutive theory of biological growth in the Föppl–von Kármán limit,” Journal of the Mechanics and Physics of Solids, vol. 57, no. 3, pp. 458–471, 2009.

17. E. K. Rodriguez, A. Hoger, and A. D. McCulloch, “Stress-dependent finite growth in soft elastic tissues,” Journal of biomechanics, vol. 27, no. 4, pp. 455–467, 1994.

18. J. V. Beira and R. Paro, “The legacy of drosophila imaginal discs,” Chromosoma, vol. 125, no. 4, pp. 573–592, 2016.

19. L. LeGoff, H. Rouault, and T. Lecuit, “A global pattern of mechanical stress polarizes cell divisions and cell shape in the growing drosophila wing disc,” Development, vol. 140, no. 19, pp. 4051–4059, 2013.

20. Y. Mao, A. L. Tournier, A. Hoppe, L. Kester, B. J. Thompson, and N. Tapon, “Differential proliferation rates generate patterns of mechanical tension that orient tissue growth,” The EMBO journal, vol. 32, no. 21, pp. 2790–2803, 2013.

21. C. Rauskolb, S. Sun, G. Sun, Y. Pan, and K. D. Irvine, “Cytoskeletal tension inhibits hippo signaling through an ajuba-warts complex,” Cell, vol. 158, no. 1, pp. 143–156, 2014.

22. J. C. Pastor-Pareja and T. Xu, “Shaping cells and organs in drosophila by opposing roles of fat body-secreted collagen iv and perlecan,” Developmental cell, vol. 21, no. 2, pp. 245–256, 2011.

23. L. Sui, G. O. Pflugfelder, and J. Shen, “The dorsocross t-box transcription factors promote tissue morphogenesis in the drosophila wing imaginal disc,” Development, vol. 139, no. 15, pp. 2773–2782, 2012.

24. L. Sui, S. Alt, M. Weigert, N. Dye, S. Eaton, F. Jug, E. W. Myers, F. Jülicher, G. Salbreux, and C. Dahmann, “Differential lateral and basal tension drive folding of drosophila wing discs through two distinct mechanisms,” Nature communications, vol. 9, no. 1, pp. 1–13, 2018.

25. W. Ramos-Lewis and A. Page-McCaw, “Basement membrane mechanics shape development: Lessons from the fly,” Matrix Biology, vol. 75, pp. 72–81, 2019.

26. A. Page-McCaw, A. J. Ewald, and Z. Werb, “Matrix metalloproteinases and the regulation of tissue remodelling,” Nature reviews Molecular cell biology, vol. 8, no. 3, pp. 221–233, 2007.

27. C. Bergantiños, M. Corominas, and F. Serras, “Cell death-induced regeneration in wing imaginal discs requires jnk signalling,” Development, vol. 137, no. 7, pp. 1169–1179, 2010.

28. A. Goriely, The mathematics and mechanics of biological growth, vol. 45. Springer, 2017.

29. E. L. L.D Landau, “Théorie de l’élasticité,” 1967.

30. E. Ruocco and M. Fraldi, “Critical behavior of flat and stiffened shell structures through different kinematical models: A comparative investigation,” Thin-walled structures, vol. 60, pp. 205–215, 2012.

31. B. Audoly and Y. Pomeau, Elasticity and Geometry. Oxford University Press, 2010.

32. I. Bock and J. Jarušek, “Dynamic contact problem for a von kármán–donnell shell,” ZAMM-Journal of Applied Mathematics and Mechanics/Zeitschrift für Angewandte Mathematik und Mechanik, vol. 93, no. 10-11, pp. 733–744, 2013.

33. P. G. Ciarlet and B. Miara, “Justification of the two-dimensional equations of a linearly elastic shallow shell,” Communications on pure and applied mathematics, vol. 45, no. 3, pp. 327–360, 1992.

34. D.-G. Zhang and Y.-H. Zhou, “A theoretical analysis of fgm thin plates based on physical neutral surface,” Computational Materials Science, vol. 44, no. 2, pp. 716–720, 2008.

35. L. O. Larbi, A. Kaci, M. S. A. Houari, and A. Tounsi, “An efficient shear deformation beam theory based on neutral surface position for bending and free vibration of functionally graded beams#,” Mechanics Based Design of Structures and Machines, vol. 41, no. 4, pp. 421–433, 2013.

36. A. Cutolo, V. Mallardo, M. Fraldi, and E. Ruocco, “Third-order nonlocal elasticity in buckling and vibration of functionally graded nanoplates on winkler-pasternak media,” Annals of Solid and Structural Mechanics, vol. 12, no. 1, pp. 141–154, 2020.

37. M. Kozlov and M. Winterhalter, “Elastic moduli for strongly curved monoplayers. position of the neutral surface,” Journal de Physique II, vol. 1, no. 9, pp. 1077–1084, 1991.

38. L. LeGoff, H. Rouault, and T. Lecuit, “A global pattern of mechanical stress polarizes cell divisions and cell shape in the growing drosophila wing disc,” Development, vol. 140, no. 19, pp. 4051–4059, 2013.

39. A. Boudaoud, P. Patrício, Y. Couder, and M. Ben Amar, “Dynamics of singularities in a constrained elastic plate,” Nature, vol. 407, no. 6805, pp. 718–720, 2000.

40. P. Bayly, R. Okamoto, G. Xu, Y. Shi, and L. Taber, “A cortical folding model incorporating stress-dependent growth explains gyral wavelengths and stress patterns in the developing brain,” Physical biology, vol. 10, no. 1, p. 016005, 2013.

41. M. B. Amar and A. Bordner, “Mimicking cortex convolutions through the wrinkling of growing soft bilayers,” Journal of Elasticity, vol. 129, no. 1, pp. 213–238, 2017.

42. D. Ambrosi, M. Ben Amar, C. J. Cyron, A. DeSimone, A. Goriely, J. D. Humphrey, and E. Kuhl, “Growth and remodelling of living tissues: perspectives, challenges and opportunities,” Journal of the Royal Society Interface, vol. 16, no. 157, p. 20190233, 2019.

43. J. Weickenmeier, C. Fischer, D. Carter, E. Kuhl, and A. Goriely, “Dimensional, geometrical, and physical constraints in skull growth,” Physical review letters, vol. 118, no. 24, p. 248101, 2017.

44. K. Alessandri, B. R. Sarangi, V. V. Gurchenkov, B. Sinha, T. R. Kießling, L. Fetler, F. Rico, S. Scheuring, C. Lamaze, A. Simon, et al., “Cellular capsules as a tool for multicellular spheroid production and for investigating the mechanics of tumor progression in vitro,” Proceedings of the National Academy of Sciences, vol. 110, no. 37, pp. 14843–14848, 2013.

45. A. Kiran, N. Kumar, and V. Mehandia, “Distinct modes of tissue expansion in free versus earlier-confined boundaries for more physiological modeling of wound healing, cancer metastasis, and tissue formation,” ACS omega, vol. 6, no. 17, pp. 11209–11222, 2021.

46. Y. Inoue, I. Tateo, and T. Adachi, “Epithelial tissue folding pattern in confined geometry,” Biomechanics and modeling in mechanobiology, pp. 1–8, 2019.

47. E. Ruocco and M. Fraldi, “An analytical model for the buckling of plates under mixed boundary conditions,” Engineering structures, vol. 38, pp. 78–88, 2012.

48. R. Farhadifar, J.-C. Röper, B. Aigouy, S. Eaton, and F. Jülicher, “The influence of cell mechanics, cell-cell interactions, and proliferation on epithelial packing,” Current Biology, vol. 17, no. 24, pp. 2095–2104, 2007.

49. D. B. Staple, R. Farhadifar, J.-C. Röper, B. Aigouy, S. Eaton, and F. Jülicher, “Mechanics and remodelling of cell packings in epithelia,” The European Physical Journal E, vol. 33, no. 2, pp. 117–127, 2010.

50. F. Graner and J. A. Glazier, “Simulation of biological cell sorting using a two-dimensional extended potts model,” Physical review letters, vol. 69, no. 13, p. 2013, 1992.

51. D. L. Weaire and S. Hutzler, The physics of foams. Oxford University Press, 2001.

52. C. Bielmeier, S. Alt, V. Weichselberger, M. La Fortezza, H. Harz, F. Jülicher, G. Salbreux, and A.-K. Classen, “Interface contractility between differently fated cells drives cell elimination and cyst formation,” Current Biology, vol. 26, no. 5, pp. 563–574, 2016.

53. E. Hannezo, J. Prost, and J.-F. Joanny, “Theory of epithelial sheet morphology in three dimensions,” Proceedings of the National Academy of Sciences, vol. 111, no. 1, pp. 27–32, 2014.

54. I. Heemskerk, T. Lecuit, and L. LeGoff, “Dynamic clonal analysis based on chronic in vivo imaging allows multiscale quantification of growth in the drosophila wing disc,” Development, vol. 141, no. 11, pp. 2339–2348, 2014.

55. F. Abouakil, H. Meng, M.-A. Burcklen, H. Rigneault, F. Galland, and L. LeGoff, “An adaptive microscope for the imaging of biological surfaces,” Light: Science & Applications, vol. 10, no. 1, pp. 1–12, 2021.

